# Artificial Microbiome-Selection to Engineer Microbiomes That Confer Salt-Tolerance to Plants

**DOI:** 10.1101/081521

**Authors:** Ulrich G Mueller, Thomas E Juenger, Melissa R Kardish, Alexis L Carlson, Kathleen Burns, Joseph A Edwards, Chad C Smith, Chi-Chun Fang, David L Des Marais

## Abstract

We develop a method to artificially select for rhizosphere microbiomes that confer salt-tolerance to the model grass *Brachypodium distachyon*. We differentially propagate microbiomes within the background of a non-evolving, highly-inbred plant population, and therefore only microbiomes evolve in our experiment, but not the plants. To optimize methods, we conceptualize artificial microbiome-selection as a special case of *indirect selection*: We do not measure microbiome properties directly, but we use host performance (e.g., biomass; seed set) as an indicator to infer association with rhizosphere microbiomes that confer salt-tolerance to a plant. We previously called this indirect-selection scheme *host-mediated indirect selection on microbiomes* (Mueller & Sachs 2015). Our methods aim to maximize evolutionary changes due to differential microbiome-propagation, while minimizing some (but not all) ecological processes affecting microbiome composition. Specifically, our methods aim to maximize microbiome perpetuation between selection-cycles and maximize response to artificial microbiome-selection by (a) controlling microbiome assembly when inoculating seeds at the beginning of each selection cycle; (b) using low-carbon soil to enhance host-control mediated by carbon secretions of plants during initial microbiome assembly and subsequent microbiome persistence; (c) fractionating microbiomes before transfer between plants to perpetuate and select only on bacterial and viral (but not fungal) microbiome components; and (d) ramping of salt-stress between selection-cycles to minimize the chance of over-stressing plants. Our selection protocol generates microbiomes that enhance plant fitness after only 1-3 rounds of artificial selection on rhizosphere microbiomes. Relative to fallow-soil control treatments, artificially-selected microbiomes increase plant fitness by 75% under sodium-sulfate stress, and by 38% under aluminum-sulfate stress. Relative to null control treatments, artificially-selected microbiomes increase plant fitness by 13% under sodium-sulfate stress, and by 12% under aluminum-sulfate stress. When testing microbiomes after nine rounds of differential microbiome propagation, the effect of bacterial microbiomes selected to confer tolerance to sodium-sulfate stress appears specific (these microbiomes do not confer tolerance to aluminum-sulfate stress), but the effect of microbiomes selected to confer tolerance to aluminum-sulfate stress appears non-specific (selected microbiomes ameliorate both sodium- and aluminum-sulfate stresses). Complementary metagenomic analyses of the artificially selected microbiomes will help elucidate metabolic properties of microbiomes that confer specific versus non-specific salt-tolerance to plants.

## INTRODUCTION

A challenge in plant-microbiome research is engineering of microbiomes with specific and lasting beneficial effects on plants. These difficulties of microbiome engineering derive from several interrelated factors, including transitions in microbiome function during plant ontogeny, and the sheer complexity of microbiome communities, such as the hyperdiverse rhizosphere or phyllosphere microbiomes containing countless fungal, bacterial, and viral components (Bulgarelli *et al* 2013; Pfeiffer *et al* 2013; Roossinck 2015). Even when beneficial microbiomes can be assembled experimentally to generate specific microbiome functions that benefit a plant, microbiomes are often ecologically unstable and undergo turnover (i.e., microbiome communities change over time), for example when new microbes immigrate into microbiomes, when beneficial microbes are lost from microbiomes, or when beneficial microbes evolve under microbe-microbe competition new properties that are detrimental to a host plant.

One strategy to engineer sustainable beneficial microbiome-function uses repeated cycles of differential microbiome-propagation to perpetuate between hosts only those microbiomes that have the most desired fitness effects on a host (Figure 1). Such differential propagation of microbiomes between hosts can therefore artificially selects for microbiome components that best mediate, for example, stresses that impact host fitness (Swenson *et al* 2000; Williams & Lenton 2007; Mueller & Sachs 2015). Only two published studies have used this approach so far (Swenson *et al* 2000; Panke-Buisse *et al* 2015). Both studies tried to select on rhizosphere microbiomes of the plant *Arabidopsis thaliana*, and both studies needed more than 10 selection-cycles to generate an overall weak and highly variable phenotypic response to microbiome-selection (e.g., increase in above-ground biomass of plants by about 10%; Swenson *et al* 2000). We here improve this differential microbiome-propagation method to artificially select for rhizosphere microbiomes that confer salt-tolerance to the model grass *Brachypodium distachyon* (Figures 1 & 2). Specifically, we build on the original methods of Swenson *et al* (2000) by modifying experimental steps that are critical to improve (i) microbiome perpetuation and (ii) response to artificial microbiome-selection. These experimental improvements include controlling microbiome assembly when inoculating seeds; low-carbon soil to enhance host-control exerted by seedlings during initial microbiome assembly and early plant growth; harvesting and perpetuating microbiomes that are in close physical contact with plants; short-cycling of microbiome-generations to select for microbiomes that benefit seedling growth; microbiome-fractionation to eliminate possible transfer of pathogenic fungi; ramping of salt-stress between selection-cycles to minimize the chance of either under-stressing or over-stressing plants.

**Figure 1.**
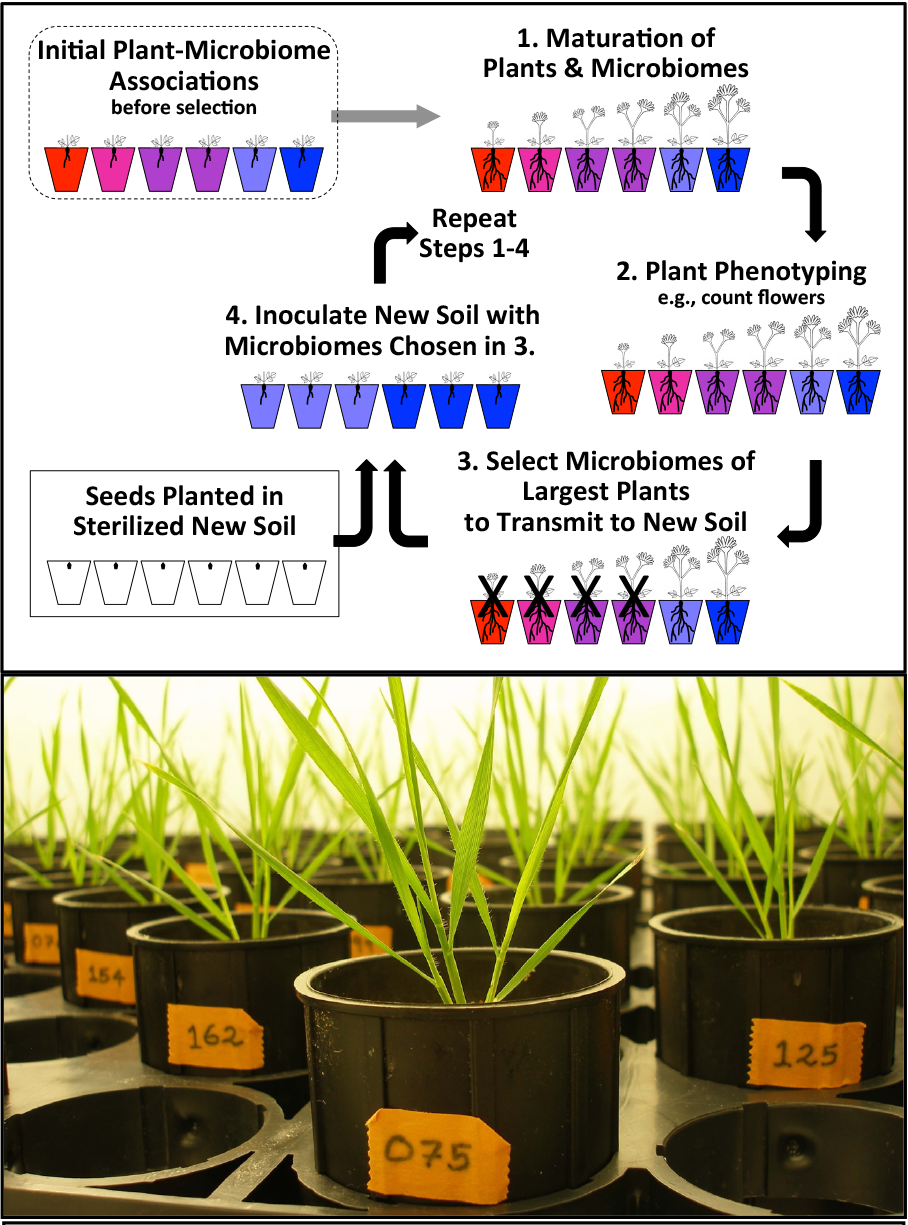
**Top:** Differential microbiome-propagation scheme to impose artificial indirect selection on rhizosphere microbiomes (figure modified from Mueller & Sachs 2015). The host-plant does not evolve because this scheme propagates only microbiomes into sterilized soil (Step 4), whereas the host is taken each cycle from a nonevolving source (stored seeds). Both evolutionary and ecological processes alter microbiomes during each cycle, but at Steps 3&4 in each cycle, experimental protocols aim to maximize evolutionary changes stemming from differential propagation of microbiomes. **Bottom:** Experimental plants of the model grass *Brachypodium distachyon* (Poaceae) shortly before microbiome-harvest for differential microbiome-propagation to seeds of the next microbiome-generation. Photo by UGM.

**Figure 2.**
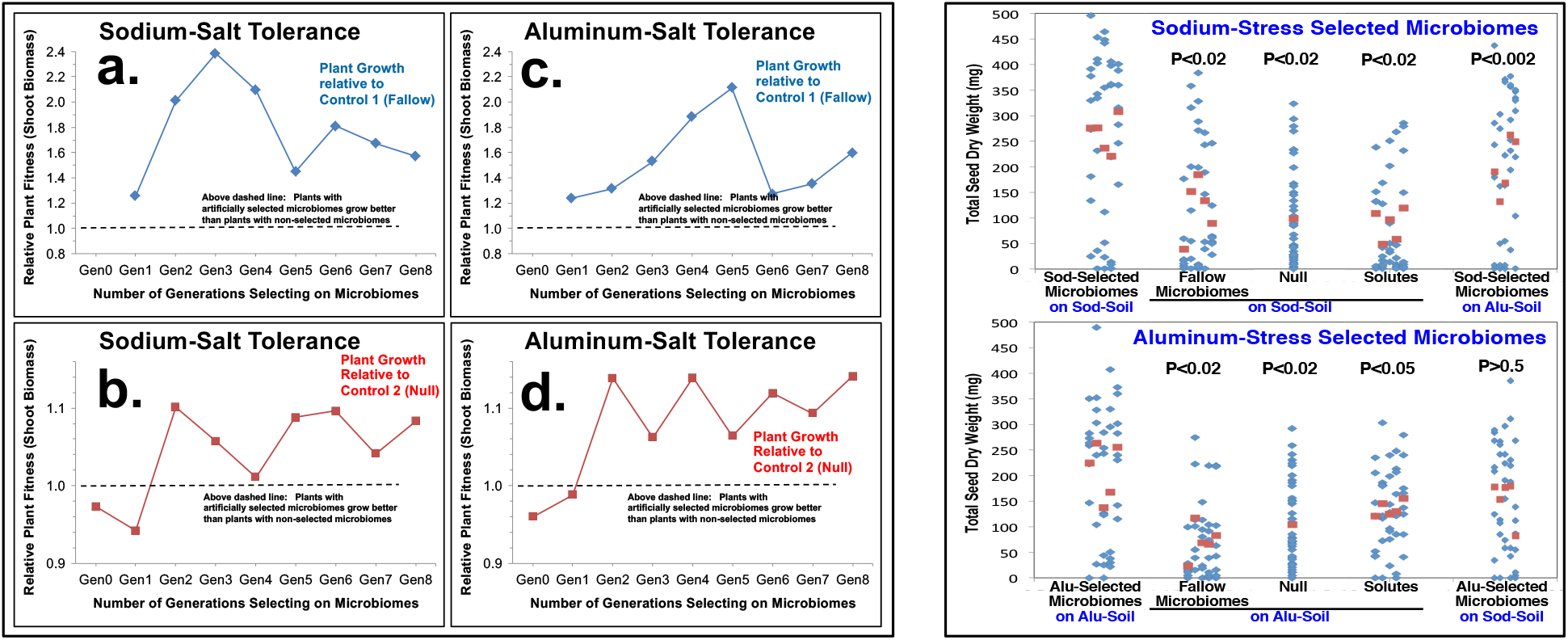
**Host-mediated artificial selection on microbiomes** to confer salt-tolerance to the grass *Brachypodium distachyon*. **Left:** Microbiomes were propagated for 8 selection cycles (generations = Gen), using the microbiome propagation scheme in Figure 1. Two salt stresses (sodium-sulfate vs. aluminum-sulfate) are contrasted, and plant fitness of selection-lines is shown relative to two non-selected control treatments. In *Fallow-Soil Microbiome-Propagation Control*, microbiomes were harvested from fallow soil (pot with no plant), then propagated to sterile fallow soil in pots of next microbiome-generation. In *Null Control*, plants did not receive a microbiome inoculum, but microbes could “rain in” from air, as in all treatments. Selection and fallow-soil treatments had 5 selection lines each, with 8 plants per line. **Right:** At the end of the experiment after a 9th selection cycle (Generation 9), plants were grown to seed for 68 days. Total seed weight is plotted in blue for individual plants (plants of same selection line are vertically above each other), the averages for each of the 5 selection lines are plotted in red. Two additional controls were added in Generation 9: *Solute-Control*, filtering out bacterial & fungal components, to test only solutes & viral microbiome components that are propagated; and *2×2 Cross-Fostering* to test for specificity of evolved microbiome function. All controls are significantly different from selection treatments, except for aluminum-selected microbiomes tested under sodium-stress (p-values are shown above each control, and are corrected using the false discovery rate for post-hoc comparisons; see Supplemental Information, Tables S3 & S4). **Conclusion:** Host-mediated artificial selection on microbiomes can generate bacterial microbiomes conferring salt-tolerance to plants; this effect is due to bacterial microbiomes and not due to solutes (plant secretions, viruses, etc.) in soil. The effect of microbiomes selected to confer tolerance to aluminum-sulfate appears non-specific (these microbiomes confer tolerance to both sodium- and aluminum-sulfate stress; p>0.5), but the effect of bacterial microbiomes selected to confer tolerance to sodium-sulfate appears specific (these microbiomes do not confer salt-tolerance to aluminum-sulfate stress; p<0.002).

### The Logic of Host-Mediated Indirect Artificial Selection on Microbiomes

To optimize microbiome-selection experiments, we found it useful to conceptualize the selection process within a host-focused quantitative-genetic framework (Mueller & Sachs 2015), rather than within a multi-level selection framework preferred by Swenson *et al* (2000; “artificial ecosystem selection”). Both frameworks capture the same processes, but a host-focused quantitative-genetic framework is more useful to identify factors that can be manipulated to increase efficacy of microbiome-selection (Mueller & Sachs 2015). First, because microbiome-selection aims to shape a fitness component of the host-plant (e.g., growth rate, stress tolerance, disease resistance), and because it is typically easier to measure plant phenotypes rather than measure microbiome properties, *selection is indirect*: Microbiomes are not measured directly, but microbiomes are evaluated indirectly by measuring host performance. Indirect selection is an established breeding technique that is often used when the indirectly-selected trait is difficult or costly to measure (Falconer & Mackay 1996), as is the case also for microbiome properties, compared to the ease of measuring a host phenotype that is dependent on microbiome properties. The efficacy of indirect selection depends on correlations between microbiome properties and host trait, and indirect microbiome-selection should therefore be more efficient if such correlations can be maximized experimentally, for example by controlling ecological priority-effects during initial microbiome assembly, or by increasing host-control over microbiome assembly and microbiome persistence. Second, because a typical host experienced a long history of evolution to monitor and manipulate its microbiomes (a process called *host control*; Sachs *et al* 2004, 2010; Kiers *et al* 2007; Berg & Smalla 2009; Bright & Bulgheresi 2010; Hillman & Goodrich-Blair 2016; Lareen *et al* 2016; Foster *et al* 2017), indirect microbiome-selection uses the host as a kind of “thermostat” to help gauge and adjust the “temperature” of its microbiomes, then propagate desired microbiomes between hosts (Mueller & Sachs 2015; Figure 1). Based on theory (Foster & Wenseleers 2006; Williams & Lenton 2007; Fitzpatrick 2014; Mueller & Sachs 2015), such *host-mediated indirect* selection on microbiomes can be easier than direct selection on microbiomes, particularly with hosts that exert strong host-control over assembly and stability of their microbiomes (Scheuring & Yu 2012; Coyte *et al* 2015; Mueller & Sachs 2015). Third, as in Swenson *et al* (2000), we used an inbred strain of a plant-host (here, *B. distachyon* genotype Bd3-1) to ensure constancy of the host-genetic environment within and between selection-cycles, such that microbiomes evolve in a specific plant-genotype background; we have previously called this method *one-sided selection* (Mueller & Sachs 2015) because only the microbiomes change between propagation cycles, whereas the host is taken each cycle from stored, inbred stock (Figure 1). The host therefore cannot evolve and microbiome-selection proceeds within the background of a genetically constant and homogenous host.

### Definition of Microbiome Engineering & Microbiome Selection

Microbiome engineering by means of differential microbiome propagation (Figure 1) alters microbiomes through both ecological and evolutionary processes. Ecological processes include changes in community diversity, relative species abundances, or structure of microbe-microbe or microbe-plant interaction networks. Evolutionary processes include extinction of specific microbiome members; allele frequency changes, mutation, or gene transfer between microbes; and differential persistence of microbiome components when differentially propagating microbiomes at each selection cycle. These processes can be interdependent (e.g., in the case of eco-evolutionary feedbacks; Strauss 2014; Theis *et al* 2016; De Meester *et al* 2019), and some processes can be called either ecological or evolutionary (e.g., loss of a microbe from a microbiome community can be viewed as evolutionary extinction or as a result of ecological competition), but for the design of an efficient microbiome-selection protocol, it is useful to think about ecological processes separately from evolutionary processes. Microbiome-selection protocols aim to maximize evolutionary changes stemming from differential microbiomes propagation (Steps 3&4 in Figure 1), for example by optimizing microbiome inheritance during microbiome transfers between hosts, or by optimizing microbiome re-assembly after such transfers (e.g., by facilitating ecological priority effects at initial host inoculation). Although both evolutionary and ecological processes alter microbiomes during each propagation cycle (Figure 1), as a shorthand, we refer here to the combined evolutionary and ecological changes resulting from host-mediated indirect artificial selection on microbiomes as ‘microbiome response’ due to ‘microbiome selection’.

## METHODS

We describe all experimental steps and analyses in great detail in the Supplemental Information, and summarize here the basic experimental approach:

### Maximizing microbiome perpetuation

To select for microbiomes that confer salt-tolerance to plants, we used a differential host-microbiome co-propagation scheme as described in Swenson *et al* (2000), Mueller *et al* (2005), and Mueller & Sachs (2015). Because both evolutionary and ecological processes alter microbiomes during and between selection-cycles (i.e., microbiome-generations), we designed a protocol that improved on these earlier selection-schemes by (i) maximizing evolutionary microbiome changes stemming from differential propagation of whole microbiomes occurring at Step 3 in Figure 1, while (ii) minimizing some, but not all, ecological microbiome changes that can occur at any of the steps in the selection cycle (e.g., we tried to minimize uncontrolled microbe-community turnover). In essence, our protocol aimed to maximize microbiome-perpetuation (i.e., maximize microbiome-inheritance of key microbes) and thus enhance efficacy of artificial indirect selection on microbiomes. To increase microbiome-inheritance, we added protocol steps of established techniques, most importantly (i) facilitation of ecological priority-effects during initial microbiome assembly (Fierer *et al* 2012; Scheuring & Yu 2012), thus increasing microbiome-inheritance by controlling in each selection-cycle the initial recruitment of symbiotic bacteria into rhizosphere microbiomes of seedlings; and (ii) low-carbon soil to enhance carbon-dependent host-control of microbiome-assembly and microbiome-persistence (Bais *et al* 2006; Bulgarelli *et al* 2013; Mueller & Sachs 2015; Coyte *et al* 2015; Tkacz & Poole 2015). Theory predicts that any experimental steps increasing fidelity of microbiome-perpetuation from mother-microbiome to offspring-microbiome should increase the efficacy of host-mediated microbiome selection (Mueller & Sachs 2015; Zeng *et al* 2017).

### Maximizing microbiome heritability

In each microbiome-propagation cycle (microbiome generation), we inoculated surface-sterilized seeds taken from non-evolving stock (inbred strain Bd3-1 of the grass *Brachypodium distachyon*; Vogel *et al* 2006; Garvin *et al* 2008; Vogel & Bragg 2009; Brkljacic *et al* 2011), using rhizosphere bacteria harvested from roots of those plants within each selection line that exhibited the greatest above-ground biomass (Figure 1). Microbiome-selection within the genetic background of an invariant (i.e., highly inbred) plant genotype increases *microbiome heritability*, defined here as the proportion of overall variation in plant phenotype that can be attributed to differences in microbiomes between plants. By keeping plant genotype invariant, microbiome heritability increases because a greater proportion of the overall plant-phenotypic variation in a selection line can be attributed to differences in microbiome-association. This increases an experimenter’s ability to identify association with a beneficial microbiome (Swenson *et al* 2000). In essence, microbiome selection within the background of a single, homogenous plant-genotype increases reliability of the host-phenotype as indicator of microbiome effects, and thus should increase the efficiency of indirect selection on microbiomes.

### Harvesting rhizosphere microbiomes & selection scheme

Each selection line consisted of a population of eight replicate plants, and each selection treatment had five replicate selection lines (i.e., 40 plants total per treatment). To phenotype plants on the day of microbiome harvesting, we judged above-ground growth visually by placing all eight plants of the same selection line in ascending order next to each other (Figure S3), then choosing the two largest plants for microbiome harvest. For all plants, we cut plants at soil level, then stored the above-ground portion in an envelope for drying and subsequent weighing. For each plant chosen for microbiome harvest, we extracted the entire root system from the soil, then harvested rhizosphere microbiomes immediately to minimize microbiome changes in the absence of plant-control in the rhizosphere. Entire root structures could be extracted whole because of a granular soil texture (Profile Porous Ceramic soil), with some loss of fine roots. Because we were interested in harvesting microbiomes that were in close physical association with roots, we discarded any soil adhering loosely to roots, leaving a root system with few firmly attached soil particles. We combined the root systems from the two best-growing plants of the same selection line, and harvested their mixed rhizophere microbiomes by immersing and gently shaking the roots in the same salt-nutrient buffer that we used also to hydrate soils (details in Supplemental Information). Combining root systems from the two best-growing plants generated a so-called *mixed microbiome* harvested from two ‘mother rhizospheres’, which we then transferred within the same selection line to all eight ‘offspring plants’ (i.e., germinating seeds) of the next microbiome generation (Figure 1).

### Microbiome fractionation with size-selecting filters before microbiome propagation

To simplify metagenomic analyses from propagated microbiomes, we used size-selecting filters (Bakken & Olsen 1987; Mueller & Sachs 2015) to filter microbiomes harvested from rhizospheres of ‘mother-plants’, thus capturing only bacteria (and possibly also viruses) for microbiome propagation to the next microbiome-generation, but eliminating from propagation between microbiome-generations any larger-celled soil-organisms with filters (i.e., we excluded filamentous fungi, protozoa, algae, mites, nematodes, etc.). This fractionation step distinguishes our methods from those of Swenson *et al* (2000) and Panke-Buisse *et al* (2015), both of which used ‘whole-community’ propagation to transfer between generations all organism living in soil, including the fungi, protozoa, algae, and multicellular organisms that were excluded from propagation through size-selecting filtering in our experiment.

### Salt-stress treatments and experimental contrasts

Using different selection-lines, we selected for beneficial microbiomes conferring salt-tolerance to either sodium-sulfate [Na_2_SO_4_] or to aluminum-sulfate [Al_2_(SO_4_)_3_]. Such an experimental contrast involving two main treatments (i.e., the two salt stresses) enables an experimenter to compare evolving microbiomes using metagenomic time-series analyses, as well as to identify candidate microbes (indicator taxa) and microbial consortia that differ between sodium- and aluminum treatments and that may therefore confer specific salt-tolerance to plants.

### Control treatments

To evaluate the effects of selection treatments, we included two non-selection control treatments. In *Fallow-Soil Microbiome-Propagation Control*, we harvested microbiomes from fallow soil (no plant growing in a pot; microbiomes are harvested from root-free soil, collected at the same depth of roots in pots with plants), then propagated the harvested microbiomes to sterile fallow soil of the next microbiome-generation. In *Null Control*, we did not inoculate germinating seeds with any microbiomes, but microbes could enter soil from air (i.e., microbial “rain”), as was the case also for all other treatments.

### Number of selection cycles (microbiome generations)

Our complete experiment involved one baseline Generation (Generation 0, Table S1a) to establish initial microbiomes in replicate pots; 8 rounds of microbiome-selection (i.e., differential microbiome-propagation) (Generations 1-8, Table S1b); and one final, ninth round (Generation 9, Table S1c) to evaluate the effects of the engineered microbiomes on flower-production and seed-set, for a total of 10 microbiome-generations.

### Ramping of salt stress

We increased salt stresses gradually during the selection experiment, by (i) increasing between generations the molarity of the water used to hydrate dry soil before soil-sterilization and planting (Table S2); and (ii) increasing correspondingly between generations also the molarity of the water used to hydrate pots at regular intervals (Table S3). Over the 10 generations, sodium-sulfate molarity in the sodium-stress treatments increased from 20 millimolar (mM) to 60 mM, and aluminum-sulfate molarity in the aluminum-stress treatments increased from 0.02 mM to 1.5 mM (Tables S2 & S3). The salt stresses of the baseline generation were chosen because, in pilot experiments, these stresses caused minimal delays in germination and growth compared to unstressed control plants. The logic of ramping and adjusting salt-stresses stepwise between selection cycles are explained further in the Supplemental Information.

## RESULTS

### Artificially-Selected Microbiomes Confer Increased Salt-Tolerance to Plants

Figure 2a-d shows the changes in relative plant-fitness (above-ground dry biomass) during 8 rounds of differential microbiome propagation. Relative to Fallow-Soil Control and Null Control treatments, selected microbiomes confer increased salt-tolerance to plants after only 1-3 selection-cycles, for both microbiomes selected to confer tolerance to sodium-stress (Figure 2a&b) or to aluminum-stress (Figure 2c&d). Relative to Fallow-Soil Controls, artificially selected microbiomes increase plant fitness by 75% under sodium-sulfate stress (p<0.001), and by 38% under aluminum-sulfate stress (p<0.001). Relative to Null Control plants, artificially selected microbiomes increase plant fitness by 13% under sodium-sulfate stress, and by 12% under aluminum-sulfate stress. Although repeated rounds of differential microbiome-propagation improved plant fitness between successive microbiome-generations (particularly in the Null Controls; Figure 2b&d), interactions between treatment and generation were not statistically significant (see Supplemental Information). This implies that fitness-enhancing effects of microbiomes from selection-lines therefore were realized after one or a few rounds of differential microbiome-propagation (e.g., Figure 2b&d), and there was insufficient statistical support that, under the gradually increasing salt stress, any additional rounds further resulted in greater plant biomass of selection-lines relative to control-lines. However, because plants were exposed to greater salt stresses in later selection cycles (see ramping of salt stress, Table S2), the selected microbiomes must have had correspondingly greater beneficial effects on the stressed plants. Selected microbiomes of later generations therefore helped plants tolerate greater salt stresses to allow plants grow the same biomass as plants in somewhat earlier generations experiencing somewhat lesser salt stress.

The phenotypic effect on plants due to the evolving microbiomes fluctuated during the eight rounds of differential microbiome propagation (Figures 2a-d; Table S1). Such fluctuations can occur in typical artificial selection experiments (e.g., chapters 10, 12 & 18 in Garland & Rose 2009), but fluctuations may be more pronounced when artificially selecting on microbiomes (Blouin *et al* 2015) because additional factors can contribute to between-generation fluctuations. Specifically, across the eight selection cycles in our experiment, the observed fluctuations could have been due to (i) uncontrolled humidity changes and correlated humidity-dependent water-needs of plants (humidity was not controlled in our growth chamber), consequently changing the effective salt-stresses; (ii) the strong ramping of salt-stress during the first five selection-cycles, possibly resulting in excessively stressed plants in Generations 4 & 5 (see discussion in Supplemental Information; Tables S2 & S3); (iii) random microbiome changes (“microbiome drift”) and consequently random microbe-microbe interactions; or (iv) other such uncontrolled factors. The fluctuations in plant fitness are most prominent during the first five selection-cycles (Figure 2a-d) when we increased salt-stress 2-fold to 5-fold between each generation and when humidity varied most (see Supplemental Information), whereas fluctuations were less pronounced during the last three generations when we changed salt-stress only minimally and humidity was relatively stable (Tables S2 & S3). These observations seem consistent with known responses of *B. distachyon* to environmental stresses (De Marais & Juenger 2011) predicting that artificial selection on microbiomes conferring salt-tolerance to plants should be most efficient under experimental conditions that rigorously control soil moisture, salt-stress, humidity, and plant transpiration.

### Effect of Artificially Selected Microbiomes on Seed Production

In the last microbiome-generation after a ninth microbiome-selection cycle (Generation 9), we grew plants for 68 days to quantify the effect of our artificially-selected microbiomes on seed production. We also added two control treatments, *Solute-Transfer Control* (*Solute Control*) and *Cross-Fostering*, to help elucidate some of the mechanisms underlying the salt-tolerance-conferring effects of selected microbiomes on seed production. In the Solute Control treatments, we eliminated with 0.2μm filters live cells from the harvested microbiomes in the selection lines, to test the growth-enhancing effects of root secretions and viruses that may be co-harvested and co-transferred with bacterial microbiomes in the selection-lines. Plants receiving these bacterial-free filtered solutes (i) had significantly poorer seed-production compared to plants that received these same solutes together with the live bacterial microbiomes (p<0.02 for sodium-stress treatment; p<0.05 for aluminum-stress treatment; Tables S3 & S4); and (ii) had seed production comparable to plants from Null Control treatments (p>0.7 for sodium-stress treatment; p>0.25 for aluminum-stress treatment; Tables S3 & S4) (Figure 2 right). These findings indicate that any plant secretions or viruses co-harvested with bacterial microbiomes did not account for the effects of the selected microbiomes conferring salt-tolerance to plants, and that any co-harvested solutes (e.g., root secretions) and viruses appear to affect plant growth like Null Control treatments.

### Specificity Test by Crossing Evolved SOD- and ALU-Microbiomes with SOD- and ALU-Stress

In the *Cross-Fostering* Control of the last microbiome-generation (Table S1c), we crossed harvested microbiomes from the sodium-stress (SOD) and aluminum-stress (ALU) selection-lines with the two types of salt-stress in soil, to test specificity of the salt-ameliorating effects of the microbiomes. The effect of microbiomes selected to confer tolerance to aluminum-sulfate appears non-specific (these aluminum-selected microbiomes confer tolerance to both sodium- and aluminum-sulfate stress; p>0.5), but the effect of bacterial microbiomes selected to confer tolerance to sodium-sulfate appears specific (these sodium-selected microbiomes do not confer salt-tolerance to aluminum-sulfate stress; p<0.002) (Figure 2 right, rightmost-graphs). The underlying microbial and metabolic mechanisms conferring specific and non-specific salt-tolerance are unknown, but metagenomic comparisons of the sodium-selected versus aluminum-selected microbiomes should help generate hypotheses that can be tested in future studies.

## DISCUSSION

Our study aimed to improve the differential microbiome-propagation scheme that was originally developed by Swenson *et al* (2000), then test the utility of our improved methods by artificially selecting on microbiomes to confer salt-stress tolerance to plants. Swenson’s original whole-soil community propagation scheme failed to generate consistent and sustained benefits for plant growth; specifically, Swenson’s growth enhancement due to the putatively selected communities was overall minor when averaged across all propagation-cycles (an average of about 10% growth enhancement), and highly variable between successive generations (Swenson *et al* 2000). To address these problems, we adopted in our experiment ideas from quantitative-genetics, microbial-ecology, and host-microbiome evolution to optimize experimental steps in our microbiome-propagation scheme, with the aim to improve perpetuation of beneficial microbiomes and improve microbiome assembly at the seedling stage. Specifically, our methods aimed to (a) facilitate ecological priority-effects during initial microbiome assembly (Fierer *et al* 2012; Scheuring & Yu 2012), thus increasing microbiome-inheritance by steering the initial recruitment of symbiotic bacteria into rhizosphere microbiomes of seedlings; (b) propagate microbiomes harvested from within the sphere of host-control (i.e., microbiomes in close physical proximity to roots) (Yan *et al* 2017; Shi *et al* 2016), whereas Swenson *et al* (2000) harvested also microbes from outside the sphere of host control; (c) enhance carbon-dependent host-control of microbiome-assembly and host-control of microbiome-persistence by using low-carbon soil (Bais *et al* 2006; Bulgarelli *et al* 2013; Mueller & Sachs 2015; Coyte *et al* 2015; Tkacz & Poole 2015); and (d) ramping of salt-stress between selection-cycles to minimize the chance of either under-stressing or over-stressing plant. Without additional experiments, it is not possible to say which of these experimental steps was most important to increase efficacy and response to microbiome-selection. The observation that it is possible to artificially select for microbiomes that confer drought tolerance to wheat grown in high-carbon potting soil (Metro-Mix 900, Jochum 2019) suggests that the low-carbon soil thought to be important in our experiment may not be essential for plant-mediated microbiome selection.

Our conceptual approach of *host-mediated indirect selection on microbiomes* (Mueller & Sachs 2015), combined with our experimental approach aiming to capitalize on evolved mechanisms of *host-control* (Sachs *et al* 2004; Mueller & Sachs 2015; Foster *et al* 2017), differs from other approaches that view microbiome-selection primarily as a multi-level selection process (“artificial ecosystem selection” sensu Swenson *et al* 2000; “meta-organism” selection or “hologenome” selection sensu Voss *et al* 2015, Rosenberg & Zilber-Rosenberg 2016, Theis *et al* 2016; “multi-layered” selection sensu Shapira 2016). While these multi-level views capture some aspects of host-microbiome population-biology (van Opstal & Bordenstein 2015; Theis *et al* 2016; Garcia & Kao-Kniffin 2018), we found that a multi-level selection view provides few useful insights for the design of microbiome-selection experiments. In contrast, basic principles of quantitative-genetics, microbial-ecology, and host-microbiome evolution (Coyte *et al* 2015; Mueller & Sachs 2015; Tkacz & Poole 2015; Rillig *et al* 2016; Wright *et al* 2019) and particularly the host-centric concept of *host control* (Sachs *et al* 2004; Foster *et al* 2017) allowed us to identify and adjust critical experimental steps and parameters in our differential microbiome-propagation scheme.

Compared to two other experiments of host-mediated microbiome-selection (Swenson *et al* 2000; Panke-Buisse *et al* 2015), our selection scheme appears to generate more pronounced and more stable effects on plant phenotype as a result of host-mediated microbiome-selection. Except for the initial two selection-cycles (Figure 2a-d), our selected microbiomes consistently outperformed in subsequent selection-cycles the non-selected microbiomes of the control conditions. In contrast, for example, Swenson *et al*’s experiments sometimes resulted in selected microbiomes that were outperformed by control microbiomes. Our methods may have generated more stable microbiome-effects because (a) only bacteria, but no fungi were propagated between generations (Swenson *et al* suspected fungal diseases as cause of complete devastation of plant populations in several of their selection cycles); (b) we may have conducted our experiment in a more stable greenhouse environment; and (c) we selected for microbiomes conferring specific benefits (salt-tolerance), rather than the non-specific, general-purpose beneficial microbiomes selected by Swenson *et al* and Panke-Buisse *et al*. After only 1-3 selection cycles, our selected microbiomes consistently outperformed the control microbiomes, with averages of 75% (SOD) and 38% (ALU) growth improvement relative to Fallow-Soil Controls; and 13% (SOD) and 12% (ALU) growth improvement relative to Null Controls (Figure 2a-d). Most importantly, when quantifying plant fitness by total seed production in the final Generation 9, our selected microbiomes outperformed Fallow-Soil Controls, Null Controls, and Solute Controls by 120-205% (SOD) and 55-195% (ALU) (Figure 2 right). Although we achieved these results under very controlled greenhouse conditions that are very different from outdoor conditions, this seems a remarkable enhancement of plant productivity compared to traditional plant breeding.

An interesting result is that microbiomes selected to benefit growth of plants during the early vegetative phase (biomass of 4-week-old plants, well before flowering; Figure 2 left) generated microbiomes with benefits that translated also into enhanced plant fitness during the reproductive phase by increasing seed set of 10-week-old plants (Figure 2 right). Rhizosphere microbiomes of grasses can change significantly during plant ontogeny (Edwards *et al* 2018), and therefore microbiomes selected to serve one function such as early growth may not necessarily optimize other functions such as seed set. Our observation that microbiome selection to promote early growth (Figure 2 left) also promotes increased seed set (Figure 2 right) therefore implies that (a) some of the same bacteria benefitting plants during the early vegetative phase under the tested salt stresses also benefit plants during reproductive phase in *B. distachyon*, despite possible changes in overall microbiome communities during plant ontogeny; (b) seed set is intrinsically tied to optimal early growth in *B. distachyon*, possibly by accelerating the timing of flowering; and (c) microbiome-selection experiments aiming to increase seed productivity do not necessarily have to select on seed set as a measured phenotype, but can shorten each selection cycle by selecting on other plant phenotypes, such as early vegetative growth.

Because our experiment was the first systematic attempt to improve the methods of Swenson *et al* (2000), we predict that it should be possible to further optimize protocols of differential microbiome propagation. Microbiome-selection therefore could emerge as a novel tool to engineer and elucidate microbiome-functions in controlled laboratory environments, and possibly also in those natural environments that allow control of key parameters affecting microbiome harvest, microbiome transfer, and microbiome inheritance. Such optimization of microbiome-selection should ideally be informed by metagenomic analyses of experimental contrasts (e.g., comparison of microbiomes selected to confer either sodium-stress versus aluminum-stress tolerance to plants) and by time-series analyses across microbiome-propagation cycles, to identify candidate microbes and microbial consortia important in mediating stresses.

## FUTURE RESEARCH TO IMPROVE METHODS OF MICROBIOME-SELECTION

To expand on our methods of artificial microbiome selection, we outline a series of additional experiments that should generate insights into key parameters that determine efficacy of microbiome selection. Arias-Sánchez *et al* (2019) and Xie *et al* (2019) recently summarized criteria for microbiome-selection experiments that are not host-mediated (e.g., selection on CO2 emission by a complex microbiome, in absence of a plant host), Lawson *et al* (2019) summarized protocols for engineering any kind of microbiome (e.g., by using bottom-up and top-down design criteria), and we focus here on methods of host-mediated microbiome selection that aim to improve performance of a plant host. Because host-mediated microbiome selection leverages traits that evolved to assemble and control microbiomes (so-called host control; Mueller & Sachs 2015, Foster *et al* 2017), the first four experiments explore whether factors promoting strong microbiome-control by a plant host could improve efficacy of microbiome selection:

1. *Artificial mirobiome-selection on endophytic vs. rhizosphere microbiomes*: Microbiomes internal to a host (e.g., endophytic microbes of plants) require some form of host infection, and therefore could be under greater host control than external microbiomes, such as rhizoplane or rhizosphere microbiomes. Consequently, under stresses that are mediated by host-controlled microbes, it may be easier to obtain a response to microbiome-selection when targeting selection on endosphere microbiomes versus external microbiomes. This prediction can be tested in an experiment that compares, in separate selection lines in the same experiment, the responses to microbiome selection when harvesting and propagating only endophytic microbiomes versus only rhizosphere microbiomes. This prediction may not hold for stresses that require stress-mediation by microbes in the external microbiome compartment of roots (e.g., microbes that detoxify toxins, such as aluminum, before they enter the root and then affect the plant negatively; for example microbes that chelate toxins external to the plant in the rhizosphere, Ma *et al* 2001; Aggarwal *et al* 2015); however, this prediction about a key role of host control for the efficacy of microbiome selection should hold for many other stresses that are mediated by microbes that a plant permits to enter into the endophytic compartment.
2. *Microbiome-selection in two genetic backgrounds differing in host-control*: A second approach to test for the role of host control is to compare microbiome-selection in two different host-genotypes, such as two inbred strains of the same plant species. For example, different host genotypes may recruit into symbiosis different kinds of microbes. Such differences in host-controlled microbiome recruitment could result in differences in microbiome-selection, and a microbiome artificially-selected within one host-genotype to improve one particular host trait may produce a different phenotypic effect when tested in a different host-genotype.
3. *Varying host control by varying carbon-content in soil*: A third approach to test host control is to compare efficacy of microbiome selection in low-versus high-carbon soil. Microbial growth in some soils is limited by carbon, and many plants therefore regulate their soil-microbiomes by carbon secretions (Zahar Haichar *et al* 2016; Sasse *et al* 2018). We therefore hypothesized that a low-carbon soil (like the carbon-free PPC soil in our experiment) may facilitate host control and consequently also microbiome-selection. This hypothesis remains to be tested, for example in a microbiome-selection experiment contrasting response to selection between soils with different carbon content. The observation that it is possible to artificially select for microbiomes that confer drought tolerance to wheat grown in high-carbon potting soil (Jochum 2019) suggests that low-carbon soil may not be essential for plant-mediated microbiome selection, but low-carbon soil could be a facilitating condition.
4. *Varying resource-limited host control by varying seed size*: A fourth approach to test host control could be to compare the efficacy of microbiome selection between plant species with large seeds versus small seeds (e.g., *Brachypodium* versus *Arabidopsis*), or between seedlings of the same plant species grown from small versus large seeds. A germinating seed has to allocate resources to above-ground growth to fix carbon and to below-ground growth to access nutrients and water, and seedlings growing from resource-rich large seeds therefore may be better able to allocate additional resources to manipulate microbiomes effectively, for example by root secretions. If such resource-allocation constraints exist for young seedlings, this could explain why our microbiome-selection experiment with *B. distachyon* appears to have generated stronger and faster response to microbiome selection compared to other such experiments with *Arabidopsis thaliana* (Swenson *et al* 2000; Panke-Buisse *et al* 2015). The best tests of seed-resource dependent efficacy of microbiome selection will not be comparisons between species, however, but within-species comparisons of the efficacy of microbiome selection within host populations grown from resource-rich large seeds versus resource-poor small seeds of the same plant species.
5. *Propagation of fractionated vs. whole microbiomes*: Experimental microbiome-propagation between host generations can be complete [all soil-community members are propagated between hosts, as in Swenson *et al* (2000) and Panke-Buisse *et al* (2015)] or microbiomes can be fractionated by excluding specific microbial components, as in our protocol where we propagated only organisms of bacterial or smaller sizes. We used fractionated microbiome-propagation because (a) we were more interested in elucidating contributions to host fitness of the understudied bacterial components than the fungal components (e.g., mycorrhizal fungi), and (b) fractionation simplifies analyses of the microbiome-responses to selection (e.g., bacterial microbiome components, but not necessarily fungal components, need to be analysed with metagenomic techniques). However, because fungal components and possible synergistic fungal-bacterial interactions cannot be selected on when using our fractionated microbiome-propagation scheme, we hypothesized previously (Mueller & Sachs 2015) that selection on fractionated microbiomes may show attenuated selection responses compared to selection on whole microbiomes. This can be tested in an experiment comparing the response to microbiome selection when propagating fractionated versus whole microbiomes, for example by using different size-selecting filters.
6. *Propagation of mixed vs. un-mixed microbiomes*. When propagating microbiomes to new hosts, it is possible to propagate mixed microbiomes harvested from different hosts, or only un-mixed microbiomes. Mixed vs. un-mixed propagation schemes therefore represent two principal methods of microbiome selection (Swenson *et al* 2000; Williams & Lenton 2007; Mueller & Sachs 2015; Rillig *et al* 2016). We hypothesized previously (Mueller & Sachs 2015) that mixed propagation may generate a faster response to microbiome-selection, but the respective advantages of mixed versus un-mixed propagation have yet to be tested. Mixed propagation may be superior to un-mixed propagation, for example if mixing generates novel combinations of microbes with novel beneficial effects on a host (Mueller & Sachs 2015), or may merge previously separate networks of microbes into a superior compound network (so-called community-network coalescence; Rillig *et al* 2016), or generate novel competitive interactions between microbes that increase microbiome stability (Coyte *et al* 2015).

## ACKNOWLEDGEMENTS

We thank Shane Merrell for help with greenhouse maintenance; Michael Mahometa for statistical advice; John Willis for suggesting the fallow-soil control treatment; Hannah Marti for suggesting the thermostat metaphor; John Vogel, Joey Knelman, Hannah Marti, Rong Ma, and Emma Dietrich for comments on the manuscript. We are grateful for funding from the Undergraduate Research Fellowship Program of the College of Natural Science at the University of Texas Austin (to KB), XXX (to JAE), XXX (to TEJ), XXX (to DLDM), the National Science Foundation (awards DEB0919519 & DEB1354666 to UGM), and the Stengl Endowment of the University of Texas at Austin.

## SUPPLEMENTAL INFORMATION: METHODS & RESULTS

### Protocol Outline

We used a differential host-microbiome co-propagation scheme as described in Swenson *et al* (2000) and in Mueller *et al* (2005) (Figure S1), but we added to this scheme protocol steps of established techniques, including (a) microbiome-fractionation using size-selecting filters (Bakken & Olsen 1987; Mueller & Sachs 2015); (b) ramping of stress in successive selection cycles (Garland & Rose 2009); (c) facilitation of priority effects during microbiome assembly (Fierer *et al* 2012; Scheuring & Yu 2012) by capping pots for the first 4 days of the germination stage (i.e., we used a so-called *semi-open system*; Mueller & Sachs 2015), thus controlling in each selection cycle the initial recruitment of symbiotic bacteria into rhizosphere microbiomes of seedlings; and (d) low-carbon soil to enhance carbon-dependent host-control of microbiome assembly and microbiome persistence (Bais *et al* 2006; Bulgarelli *et al* 2013; Mueller & Sachs 2015; Coyte *et al* 2015). In each microbiome-propagation cycle (‘Microbiome Generation’= Gen), we inoculated surface-sterilized seeds taken from non-evolving stock (inbred strain Bd3-1 of the grass *Brachypodium distachyon*, derived via single-seed-descent inbreeding from the source accession; Vogel *et al* 2006; Garvin *et al* 2008; Vogel & Bragg 2009; Brkljacic *et al* 2011). We chose to conduct the experiment with *B. distachyon* because it is a model for biofuel and cereal crops, including research on salt stresses and water-use efficiency (De Marais & Juenger 2011; De Marais et al 2016).

**Figure S1.**
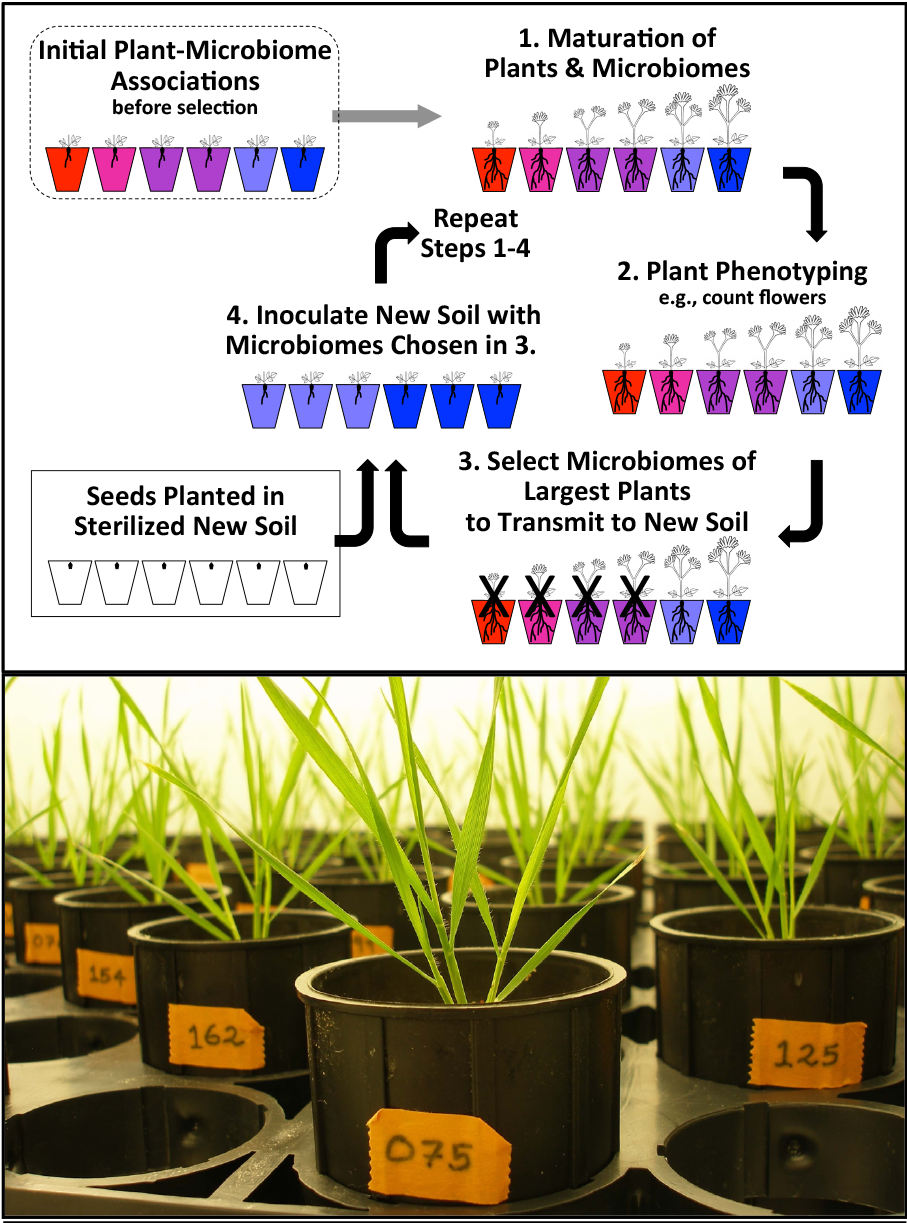
**Top:** Differential microbiome-propagation scheme to impose artificial indirect selection on rhizosphere microbiomes (figure modified from Mueller & Sachs 2015). The host-plant does not evolve because this scheme propagates only microbiomes into sterilized soil (Step 4), whereas the host is taken each cycle from a nonevolving source (stored seeds). Both evolutionary and ecological processes alter microbiomes during each cycle, but at Steps 3&4 in each cycle, experimental protocols aim to maximize evolutionary changes stemming from differential propagation of microbiomes. **Bottom:** *Brachypodium distachyon* Bd3-1 plants in our growth chamber shortly before a microbiome-harvest for differential microbiome-propagation to soils/seeds of the next microbiome-generation. Photo by UGM.

We inoculated seeds with rhizosphere bacteria harvested from roots of those plants of the previous selection cycle that exhibited the greatest above-ground biomass (Figure S1). Because the plant-host could not evolve between selection-cycles (seeds were taken from non-evolving stock), whereas microbiomes could potentially evolve due to differential microbiome propagation, our selection-scheme was *one-sided selection* (Mueller & Sachs 2015). Both evolutionary and ecological processes alter microbiomes during and between selection-cycles, but our protocol aimed to maximize evolutionary changes stemming from differential microbiome-propagation (Figure S1). To focus indirect selection on bacterial communities, we filtered the microbiomes harvested from rhizospheres, perpetuating only bacteria (and possibly also viruses) to the next generation, but eliminating from propagation between microbiome-generations any larger-celled soil-organisms with filters (i.e., we excluded fungi, protozoa, algae, mites, nematodes, etc.). This fractionation step distinguishes our methods from those of Swenson *et al* (2000) and from a replication of that study by Panke-Buisse *et al* (2015), both of which used differential ‘whole-community’ propagation to transfer between generations all organism living in soil, including the larger-celled fungi, protozoa, algae, mites, and nematodes that were excluded through size-selecting filtering in our experiment. Our complete experiment involved one baseline Generation (Generation 0, Table S1a) to establish initial microbiomes in replicate pots; 8 rounds of microbiome-selection (i.e., differential microbiome-propagation) (Generations 1-8, Table S1b); and one final ninth round of selection (Generation 9, Table S1c) to evaluate the effects of the engineered (i.e., evolved) microbiomes on flower-production and seed-set, for a total of 10 Generations. Our entire selection experiment lasted 300 days from 3. January-29. October 2015 (Table S2).

**Table S2.**
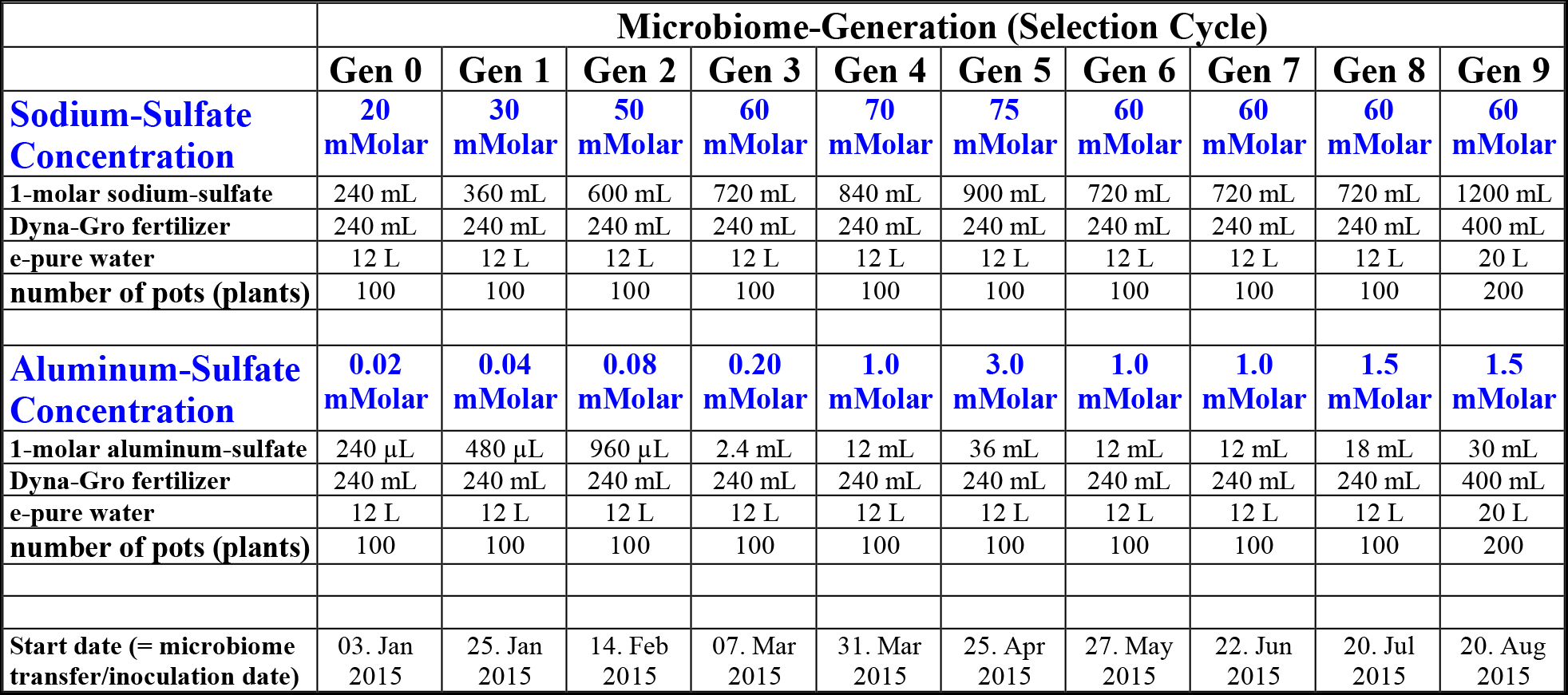
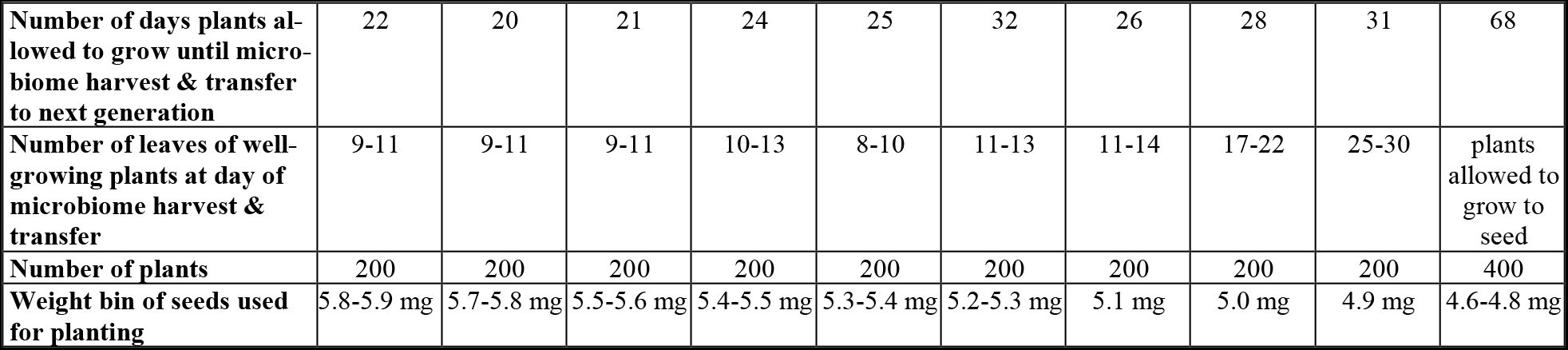
**Salt concentrations (Millimolar = mMolar) of salt-nutrient solutions used to hydrate soil** for each selection cycle (= Microbiome-Generation = Gen); the recipes to mix these solutions; and growth parameters for each Generation. In the short-cycled Generations 0-8, time was too short for plants to flower, and we quantified plant-performance by visually estimating above-ground biomass (see *Phenotyping of Plants*). In Generation 9, plants were grown for 68 days to produce seeds, and we quantified plant-performance as total seed weight per plant.

|  | **Microbiome-Generation (Selection Cycle)** |  |  |  |  |  |  |  |  |  |
| --- | --- | --- | --- | --- | --- | --- | --- | --- | --- | --- |
|  | **Gen 0** | **Gen 1** | **Gen 2** | **Gen 3** | **Gen 4** | **Gen 5** | **Gen 6** | **Gen 7** | **Gen 8** | **Gen 9** |
| **Sodium-Sulfate Concentration** | **20 mMolar** | **30 mMolar** | **50 mMolar** | **60 mMolar** | **70 mMolar** | **75 mMolar** | **60 mMolar** | **60 mMolar** | **60 mMolar** | **60 mMolar** |
| 1-molar sodium-sulfate | 240 mL | 360 mL | 600 mL | 720 mL | 840 mL | 900 mL | 720 mL | 720 mL | 720 mL | 1200 mL |
| Dyna-Gro fertilizer | 240 mL | 240 mL | 240 mL | 240 mL | 240 mL | 240 mL | 240 mL | 240 mL | 240 mL | 400 mL |
| e-pure water | 12 L | 12 L | 12 L | 12 L | 12 L | 12 L | 12 L | 12 L | 12 L | 20 L |
| **number of pots (plants)** | 100 | 100 | 100 | 100 | 100 | 100 | 100 | 100 | 100 | 200 |
| **Aluminum-Sulfate Concentration** | **0.02 mMolar** | **0.04 mMolar** | **0.08 mMolar** | **0.20 mMolar** | **1.0 mMolar** | **3.0 mMolar** | **1.0 mMolar** | **1.0 mMolar** | **1.5 mMolar** | **1.5 mMolar** |
| 1-molar aluminum-sulfate | 240 µL | 480 µL | 960 µL | 2.4 mL | 12 mL | 36 mL | 12 mL | 12 mL | 18 mL | 30 mL |
| Dyna-Gro fertilizer | 240 mL | 240 mL | 240 mL | 240 mL | 240 mL | 240 mL | 240 mL | 240 mL | 240 mL | 400 mL |
| e-pure water | 12 L | 12 L | 12 L | 12 L | 12 L | 12 L | 12 L | 12 L | 12 L | 20 L |
| **number of pots (plants)** | 100 | 100 | 100 | 100 | 100 | 100 | 100 | 100 | 100 | 200 |
| **Start date (= microbiome transfer/inoculation date)** | 03. Jan 2015 | 25. Jan 2015 | 14. Feb 2015 | 07. Mar 2015 | 31. Mar 2015 | 25. Apr 2015 | 27. May 2015 | 22. Jun 2015 | 20. Jul 2015 | 20. Aug 2015 |
| Number of days plants allowed to grow until microbiome harvest & transfer to next generation | 22 | 20 | 21 | 24 | 25 | 32 | 26 | 28 | 31 | 68 |
| Number of leaves of well-growing plants at day of microbiome harvest & transfer | 9-11 | 9-11 | 9-11 | 10-13 | 8-10 | 11-13 | 11-14 | 17-22 | 25-30 | plants allowed to grow to seed |
| Number of plants | 200 | 200 | 200 | 200 | 200 | 200 | 200 | 200 | 200 | 400 |
| Weight bin of seeds used for planting | 5.8-5.9 mg | 5.7-5.8 mg | 5.5-5.6 mg | 5.4-5.5 mg | 5.3-5.4 mg | 5.2-5.3 mg | 5.1 mg | 5.0 mg | 4.9 mg | 4.6-4.8 mg |

### Logic of Salt-Stress Ramping

We used ramping of salt-stress (Mueller & Sachs 2015) to ensure that (a) plants were neither under-stressed nor excessively over-stressed during any selection-cycle of our microbiome-selection experiment, and thus (b) facilitate that microbiomes can gradually improve under differential microbiome-propagation to confer increasingly greater salt-tolerance to plants under increasingly greater salt-stress. The experimental rationale of stress-ramping is as follows: if salt-stress is too weak, all plants grow well, any salt-stress-mediating microbiomes will make little or no difference to plants, and no microbiome-mediated variation in plant-phenotype may emerge that could be used as direct target for indirect selection on microbiomes; in contrast, if salt-stress is excessive, all plants suffer severely, and any observed variation in plant-phenotype may be due to microbiome-unrelated effects emerging under excessive stress, such that possible beneficial effects of salt-stress-mediating microbiomes are dwarfed and masked by the excessive stress. Stress-ramping is therefore an experimental trick that permits an experimenter to continuously adjust stress during a selection experiment, particularly in experimental evolution where the evolving effect sizes cannot be known *a priori* (i.e., in our experiment, it was not possible to predict *a priori* the approximate effect sizes attributable to beneficial microbiomes that could emerge as a result of multiple rounds of differential microbiome-propagation).

Table S2 shows the ramped salt-concentrations for the two salt treatments of soils in our experiment, Na_2_SO_4_ (sodium-sulfate, henceforth *SOD-soil treatment*) and Al_2_(SO_4_)_3_ (aluminum-sulfate, *ALU-soil treatment*). We chose the particular two salt stresses because sodium-cations are a problem in saline and sodic soils (e.g., Lodeyro & Carrillo 2015), and aluminum-cations are a problem because aluminum inhibits, at even minimum concentrations, plant growth in low-pH soils (Delhazie *et al* 1995; Aggarwal *et al* 2015). Our maximum sodium-salt stress of 75 mMolar salt-concentration sodium-sulfate of water used to hydrate soil and water plants during the experiment is not quite comparable to the salt stress of 500 mMolar sodium-chloride used by Priest *et al* (2014) because (a) the two experiments used different kinds of salts and (b) Priest *et al* spiked salt stress after initial growth of unstressed plants, whereas in our experiment the plants were salt-stressed already at the germination stage and at all times during each selection cycle.

A second pre-planned feature of our experimental design was to use ‘short-cycling’ in the initial selection-cycles (cycling at about 20-day intervals; plants grew to about the 9-13 leaf stage to grow sufficiently large root systems for microbiome harvest, but plants did not have sufficient time to flower), and then to increase lengths of selection-cycles gradually as plants became more stressed under the ramped salt-concentrations and plants needed more time to grow to the 9-13 leaf stage. Although we planned lengthening the duration of selection-cycles during our multi-generation experiment, we did not pre-plan at the beginning of our experiment the exact length of each selection-cycle, because the exact transfer dates were dependent also on time-constraints of the main experimenter (UGM) handling the microbiome-transfers. Because we increased salt-stress during the 10-Generation experiment (Table S2), plant growth was slower in later generations.

### Preparation of *Brachypodium distachyon* Seeds

Prior to the start of the microbiome-selection experiment, we harvested about 6000 seeds from 36 plants (*B. distachyon* strain Bd3-1; Garvin *et al* 2008; Vogel & Bragg 2009) grown simultaneously at room temperature under constant light-cycle (14h light, 10h dark) in well-homogenized, well-watered and well-fertilized greenhouse potting soil. Seeds were air-dried at room temperature for 4 months, mixed well, then weighed individually to the nearest 0.1mg to generate cohorts of seeds of equal weight (binned to within 0.1mg). To reduce within-generation phenotypic variation due to differences in seed-weight-dependent maternal effects, we used seeds of only one or two adjacent weight-bins for each generation (see last row in Table S2). We used seeds of 5.9&5.8mg weight for the initial baseline Generation 0, then we used up seeds of bins of gradually decreasing seed-weight (5.9&5.8mg, 5.8&5.7mg, 5.6&5.5mg, …), as shown in Table S2 for each microbiome-generation. All microbiome selection-cycles used seeds from this stored (non-evolving) seed-stock of Bd3-1 plants, and microbiomes were therefore selected under a so-called *one-sided selection* scheme in the single plant-genotype background Bd3-1 (only microbiomes can change between selection-cycles, but the plant-host cannot evolve; Mueller & Sachs 2015).

### Growth Chamber

For the multi-generation selection experiment, we grew plants under constant temperature (24°C) and constant light-cycle (20h light 4AM-midnight, 4h dark) in a walk-in growth chamber (model MTPS72; Conviron, Winnipeg, Canada) at the Welch Greenhouse Facility of the University of Texas at Austin. The chamber was not humidity-controlled, and chamber humidity therefore varied with outdoor humidity/rainfall and with any heating (in winter) affecting humidity of the air circulating in the Greenhouse Facility. Because of unusual rainfall in spring 2015, humidity was highest in the growth chamber during Generations 4 & 5 (Table S1 & S2), and lowest during selection-cycles 0-2 and 7-9. Unfortunately, we did not monitor exact humidity with a hygrometer in the chamber, but we recorded in a journal any days of high humidity. We grew plants on two shelves (each 120cm x 100cm) in the Conviron chamber, under fluorescent lights (Sylvania T8 fluorescent tubes spaced at 10cm, plus a center row of T2 fluorescent spiral-bulbs) generating a light-intensity of 192 μmol/m^2^/s at soil level. Except for preparation of pots and planting of seeds, we performed all experimental steps for artificial microbiome-selection in this chamber, including microbiome-harvesting from rhizospheres, microbiome-fractionation (filtering), and microbiome-transfers to surface-sterilized seeds planted in sterile soil (details below).

### Soil & Pot Preparation

We grew plants from surface-sterilized seeds, each planted individually in the center of its own D50-Deepot (5cm pot diameter, 17.8cm depth, total volume 262ml; model D16H; Stewe & Sons, Tangent, Oregon, USA) filled with autoclaved PPC soil (Profile Porous Ceramic soil, Greens-Grade™ Emerald, Natural Color; PROFILE Products LLC, Buffalo Grove, IL, USA). To permit autoclaving of soil in the Deepots prior to planting, we pressed heat-tolerant fiberglass-fill into the bottom of each pot to plug bottom-drainage holes, then compacted dry PPC soil into each pot until the soil level reached 15mm below the pot margin. Each plug consisted of a fiberglass square (9.5cm x 9.5cm) cut from an insulation-sheet (R-13 EcoTouch Insulation Roll; 38cm width; GreenGuard-certified, formaldehyde-free), then pressed firmly into the bottom of a pot. After compacting soil in all pots used for a given selection-cycle (200 pots in selection-cycles Generation 0-8; 400 pots in the final Generation 9) we carefully equalized soil levels between all pots.

According to the manufacturer’s website (www.profileevs.com/products/soil-amendments/profile-porous-ceramic-ppc), PPC soil is a calcined, non-swelling illite, non-crystalline opal mineral; it has 74% pore space, with 39% capillary (water) pores and 35% non-capillary (air) pores; pH = 5.5; cation-exchange-capacity of 33.6 mEq/100g; and a chemical composition of 74% SiO_2_, 11% Al_2_O_3_, 5% Fe2O3, and less than 5% of the remainder combining all other chemicals (e.g., CaO, MgO, K_2_O, Na_2_O, TiO_2_). We chose PPC soil for three reasons: First, PPC has a very homogeneous consistency because of its uniform particle size; soil-quantity and soil-quality are therefore easy to standardize between pots. Second, whole root systems can be easily extracted from hydrated soil with little rupture of roots. Third, because the manufacturer exposes PPC soil to high temperature (heated in a rotary kiln at 1200 degrees Fahrenheit, then de-dusted), the soil contains minimum carbon, and we believed that such low- or no-carbon soil could facilitate a plant’s ability for carbon-mediated host-control (via carbon secretions by roots; see above *Protocol Outline*) (Bais *et al* 2006; Bulgarelli *et al* 2013; Mueller & Sachs 2015; Coyte *et al* 2015) of microbiome-assembly and microbiome-stability.

### Soil Hydration & Salt-Stress Treatments

After compacting soil into each pot with a wooden dowel and equalizing soil levels between all pots used in a selection-cycle, we hydrated each pot with 94ml of a fertilizer-salt solution (recipes for solutions are listed in Table S2, and are described also below). The fertilizer concentrations in this solution was identical in each selection-cycle (i.e., we added the same amount of fertilizer to soil of each microbiome generation), but we increased salt-concentrations gradually between successive selection-cycles in order to ramp salt-stress, as shown in Table S2 for the two salt-stress treatments, Na_2_S0_4_ (decahydrate sodium-sulfate, MW=322.2g; *SOD-soil*) and Al_2_(S0_4_)_3_ (anhydrous aluminum-sulfate, MW=342.15; *ALU-soil*). We chose the particular two salt stresses because sodium is a problem in saline soils (e.g., Lodeyro & Carrillo 2015), and aluminum is a problem because aluminum inhibits, at even minimum concentrations, plant growth in low-pH soils (Delhazie *et al* 1995; Aggarwal *et al* 2015). Because of this pH-dependent growth-attenuating effect of aluminum in soil, we suspected that it may be easier to select for a microbiome conferring tolerance to aluminum salt, for example by selecting for a microbiome that increases soil pH (i.e., artificial microbiome-selection would select against acidifying bacteria in microbiomes). We therefore were able to formulate this *a priori* hypothesis on a possible pH-based mechanistic basis of a microbiome-conferred tolerance to aluminum-salt. In contrast, we did not formulate a similarly specific mechanistic hypothesis for why a microbiome could confer tolerance to sodium-salt, although a number of hypotheses have been suggested in the literature, such as changes in phytohormone concentrations influencing plant physiology, or indirect physiological effects on transpiration rates (Dodd & Pérez-Alfocea 2012). We included this second selection-regime selecting for microbiomes conferring sodium-tolerance because a dual experimental design of two soil-treatments (aluminum- and sodium-salt stress) offered two advantages: (i) we could contrast evolving microbiomes between aluminum-versus sodium-treatments to identify candidate bacterial taxa or candidate consortia that may be important in mediating microbiome-conferred salt-tolerance to plants; and (ii) we could cross selection history with selection stress in the last Generation 9 to test for possible specificities of evolved microbiomes, as explained further in *Crossing Evolved SOD- and ALU-Microbiomes with SOD- and ALU-Stress*.

The salt-concentration of the baseline Generation 0 (Table S2) was determined in a salt-gradient pilot experiment as that salt-concentration that caused a minimal, but just noticeable, delay in germination and a minimal growth-rate reduction. Because aluminum-sulfate delays germination and attenuates growth at far lower concentrations than sodium-sulfate, concentrations in the ALU-treatment (Table S2) were lower by several orders of magnitude than the concentrations of sodium-sulfate in the SOD-treatment. For ramping of salt-stress, pre-planned step-increments in salt-concentration between selection-cycles were likewise informed by our pilot experiments, which suggested increments for aluminum-sulfate concentrations of about two-to five-fold for the first few microbiome-generations, and less than two-fold increments for sodium-sulfate concentrations, with gradual decrease in step-increments in later microbiome-generations so as not to over-stress plants (Table S2). Because we had to prepare hydrated soil for the next selection-cycle about 1-2 weeks before the end of a given cycle, we had to decide salt-stress increments for the next selection-cycle well in advance, using information from relative growth of younger plants in the sodium-sulfate and the aluminum-sulfate treatments. Decisions on salt-increments between Generations therefore typically involved some informed guessing, to adjust salt concentrations for the next cycle such that plants in either treatment were projected to germinate and grow at about the same rate (i.e., we aimed for plants in either salt treatment to grow to comparable sizes in the same time during a selection cycle). With such projected equal growth between sodium- and aluminum-treatments, microbiomes could be harvested at the end of a selection cycle from plants of comparable sizes (typically 9-15 leaves at the time of microbiome harvesting) regardless of whether a plant was stressed with aluminum-sulfate or sodium-sulfate (i.e., sodium-treated plants were not behind in growth compared to aluminum-treated plants, or vice versa). A second preplanned feature of our salt-ramping design was to increase salt-stress in successive selection-cycles as long as differences in effect-sizes seemed to increase between salt- and control-treatments, but to reduce the salt-stress if differences in effect-sizes diminished or disappeared, possibly because of over-stressing the plants (see above *Logic of Salt-Stress Ramping*). This seemed to happen in Generations 4 & 5 (see Figure 1 in main text), and salt-stress was therefore reduced somewhat in the subsequent four Generations 6-9 (Table S2).

For hydration of 100 pots, we mixed, in a large carboy, 12 liter double-distilled e-pure water at a 50:1-ratio with 240ml Dyna-Gro 9-7-5 (Nutrient Solutions, Richmond, CA; www.dyna-gro.com/795.htm), plus an aliquot of 1-Molar salt solution (Table S2 lists salt-aliquots in recipes for salt-nutrient mixes) to generate the specific salt-stress planned for a particular selection-cycle. [To prepare 1-Molar ALU-salt stock, we dissolved 307.94g anhydrous aluminum-sulfate in 900ml e-pure water in a 1-liter bottle; to prepare 1-Molar SOD-salt stock, we dissolved 289.98g decahydrous sodium-sulfate in 900ml e-pure water in a 1-liter bottle; then filter-sterilized each salt solution to prepare sterile stock.] We used different carboys to prepare salt-nutrient mixes for the different salt treatments (SOD, ALU). The nutrient concentration in each mix (Table S2) was sufficient such that plants did not need additional fertilization during each selection-cycle of 20-30 days during Generations 0-8 when we quantified plant fitness as above-ground biomass production, and plants even had sufficient nutrients to flower and grow seed during the 68 days of the last Generation 9 when we quantified plant fitness as seed production. For both salt treatments, fertilizer-salt solutions had a pH = 3.75 before addition to soil, but because of the buffering capacity of PPC soil (natural pH = 5.5, see above), the hydrated soil had a pH of about 5.0-5.5 after autoclaving soils, using the pH-measurement procedure in ISO/FDIS 10390 (2005). After hydration of all pots, we immediately autoclaved all pots (to minimize the time that any live microbes in the soil could consume any of the nutrients), and we autoclaved in separate 1-liter flasks at the same time 800ml of each of the unused salt-nutrient solutions; these autoclaved salt-nutrient solutions were used later during planting, and as buffer (at half-concentration) to suspend microbiomes harvested from rhizospheres for microbiome-transfers (see *Planting & Microbiome-Harvest* below).

### Autoclaving of Soil

After hydration of soil by carefully pouring exactly 94ml of fertilizer-salt solution into a pot, we leveled and smoothed the soil-surface in a pot with the bottom of a glass (same size as interior diameter of a pot); taped to each pot a label of autoclavable label-tape (Fisherbrand™) with a pre-written pot-number (#001-100 for pots of SOD-treatment; #101-200 for pots of ALU-treatment) to the top side of each pot (Figure S1); then used pre-cut pieces of aluminum foil to cap the top and wrap the bottom of each pot to prevent possible microbial contamination during seed-stratification (see below *Planting & Stratification*). Wrapped pots were arranged vertically in large autoclave trays (67 pots per tray, 3 trays total), the trays were covered with sheets of aluminum foil, then all pots in these 3 trays were sterilized simultaneously in a large autoclave. Hydration, labeling and capping of a set of 200 pots needed typically 5-6 hours. The subsequent autoclaving procedure lasted about 10 hours overnight, starting in the evening with a first cycle of 35 minutes autoclaving (121C° temperature, 20 atm pressure) with a slow-exhaust phase lasting 90 minutes; followed by overnight exposure to high temperature in the unpressurized autoclave; followed in the morning by a second cycle of 35 minutes autoclaving with a 90-minute slow-exhaust phase. This stringent autoclaving regime was sufficient to sterilize PPC soil, because plating on PDA-medium of about 0.5g soil (n=2 SOD pots, n=2 ALU pots) taken with a sterile spatula from the interior of such autoclaved pots produced no visible microbial growth within a month of incubation at room temperature. After cooling of autoclaved pots in the foil-covered trays at room temperature for at least 16 hours, we planted seeds into the sterilized soil (one seed per pot; see below *Planting*).

### Seeds Preparation & Binning of Seeds by Weight

To have enough seeds for our 10-generation selection experiment, we first grew *Brachypodium distachyon* Bd3-1 plants under standardized light conditions (14h light, 10h dark) and room temperature in well-fertilized and well-watered greenhouse soil, harvested about 6000 seeds from these plants, then dried and stored seeds at room temperature (see above *Preparation of Brachypodium distachyon Seeds*). For our experiment, we used only long-awn seeds; that is, we discarded any short-awn seeds positioned peripherally in inflorescences (spikelet), and we discarded also any misshapen or discolored seeds. We used only long-awn seeds because these kind of seeds grow in more standardized central positions in a spikelet, because we could grasp an awn with a forceps during weighing and planting without risk of injuring a seed, and because we could plant seeds vertically into soil with only the awn protruding above the soil to reveal the exact location of a seed during later microbiome inoculation (see below *Seed Inoculation*). To weigh each seed accurately, we first removed any attached glumes to weigh only the seed with its awn. One experimenter pre-weighed each seed to bin seeds by weight to the nearest 0.1mg, then a second experimenter re-weighed all seeds in bins 4.5mg – 6.0mg again (i.e., each seed was weighed twice). To help reduce within-treatment variation in plant-phenotype (specifically here, reduce seed-weight-dependent maternal effects on plant-phenotypes, as illustrated in Figure S2), we used seeds of only a narrow weight-window for each microbiome-selection cycle. We used seeds of 5.9&5.8mg weight for the initial baseline Generation 0, then we used up seeds of bins of gradually decreasing seed-weight (5.9&5.8mg, 5.8&5.7mg, 5.6&5.5mg, …), as shown in Table S2 for each microbiome-generation.

**Figure S2.**
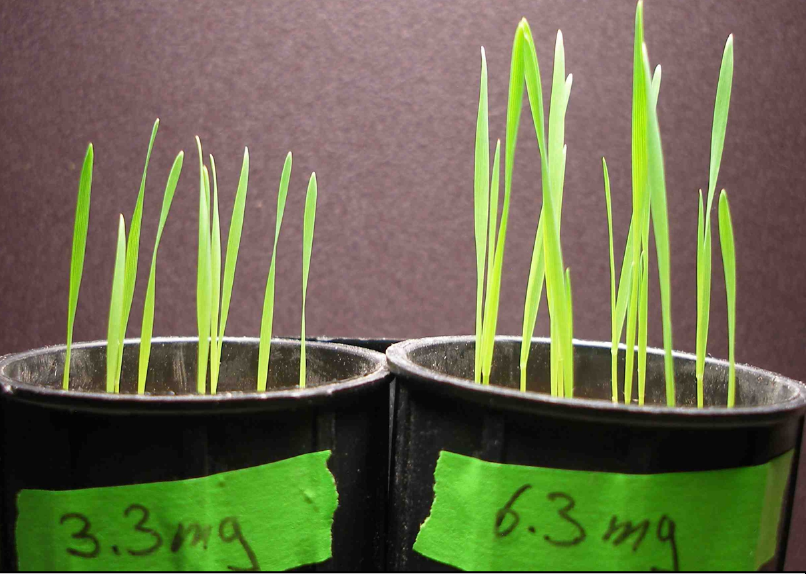
Pilot experiment illustrating growth variation of *B. distachyon* Bd3-1 plants growing under identical conditions from seeds weighing either 3.3mg or 6.3mg. The seed-weight range tested here includes about 90% of the 6000 seeds that we harvested for our microbiome-selection experiment. We used seeds of a narrow weight-window of only 0.1mg or 0.2mg for each microbiome-selection cycle (see Table S2), to help reduce within-generation and within-treatment variation in plant-phenotype (specifically here, reduce seed-weight-dependent maternal effects on phenotypes), Photo by UGM.

### Planting & Stratification

For planting of seeds in sterile soil, we first surface-sterilized Bd3-1 seeds in a laminar flow-hood by gently shaking the seeds for 8 minutes in 10% bleach [Chlorox^®^; 4ml bleach added to 36ml autoclaved e-pure water in a 50ml Falcon tube; plus 4μl Tween80-surfactant (Sigma-Aldrich, Saint Louis, MO, USA) to promote wetting of seeds], then rinsing the seeds three times to wash off bleach (succession of three 1-minute gentle shaking, each in 40ml e-pure autoclaved water in a 50-ml Falcon tube). In pilot tests, such surface-sterilized seeds placed on PDA-medium did not lead to bacterial or fungal growth. After rinsing, we blotted seeds on autoclaved filter paper, then air-dried the seeds in an open Petri dish in the flow-hood while preparing the flow-hood for planting inside the hood. To plant one seed into the center of a pot, we removed the aluminum-foil lid from a pot inside the flow-hood, pushed a narrow hole into the center of the soil with a flame-sterilized fine-tipped forceps (#5 forceps), then inserted a seed into that hole such that the seed was positioned vertically in the soil and only the awn was protruding above the soil (i.e., the pointed tip of a seed was just below the soil surface). Because seeds used for a selection cycle had been binned to within 0.1mg weight (i.e., all seeds were of same size for each selection cycle), seeds were therefore planted at the same depth (to within about 0.5-1.0 mm identical depth), and any differences in initial germination rate (i.e., appearance of the shoot at soil surface) was unlikely due to differences in planting depth between seeds. To solidify the soil around each seed, we applied 4ml autoclaved salt-nutrient solution (same concentration that was used to hydrate soil in a given selection-cycle; Table S2) with a 5ml pipette to flush soil into the hole and completely cover each seed (excepting the awn protruding vertically above the soil surface). We covered each pot with a translucent, ethanol-sterilized lid (inverted Mini Clear Plastic Bowl 40ct; Party City, Rockaway, NJ, USA). The lids prevented entry of airborne microbes into each pot, but did not seal pots completely and permitted some gas exchange. Each lid measured 5.7cm diameter x 3.8cm height, and fit snugly only each pot such that a series of 50 capped pots could be kept in a rack (D50T rack, see above) without the lids interfering with each other. We placed each rack of 50 capped pots into its own ethanol-sterilized plastic tub (Jumbo Box; Container Store, Coppell, TX), covered the box with a boxes plastic lid, then sealed the spaces at the side of each lid by wrapping lid&tub with a 2-meter-long strip of 10-cm-wide Parafilm to prevent entry of contaminants during subsequent cold-storage for stratification of seeds. We moved each tub into cold-storage immediately after completing the planting of 50 pots. For stratification, we stored the tubs with planted seeds in a 5°C cold-room for about 5 days (range 4-10 days, the duration differing slightly between selection-cycles because of scheduling-constraints affecting planting). Planting of a set of 200 seeds (4 racks of 50 pots each) using the above methods needed typically 4.5-5.5 hours.

### Preparations for Microbiome-Harvesting

To prepare salt-nutrient buffer-solution for microbiome harvesting, we used the autoclaved salt-nutrient solution that we had prepared for hydration of soil for a particular selection-cycle (Table S2), then diluted the solution to half-concentration by addition of an equal volume of autoclaved e-pure water. We decided to use for microbiome-harvesting the salt-nutrient solution at half-concentration, because we were concerned that the full-concentration may have too high osmolarity compared to the osmolarity that may exist in the soil after weeks of root- and microbiome-growth in the soil; this dilution precaution may not have been necessary, and it may be possible to propagate microbiomes even when using full-concentration of salt-nutrient solution. Aliquots of 45ml of the sterile, half-concentration salt-nutrient buffer were added to 50ml Falcon tubes in a laminar-flow hood, and these tubes were then pre-labeled with relevant information (SOD vs ALU treatment; Generation #; date of microbiome-harvest) to save time on the actual day of microbiome-harvest. To sterilize microfilters needed for fractionation of harvested microbiomes (2μm Whatman™ filters; model Puradisc 25 GD2 Syringe Filter, 25mm diameter; Whatman PLC, United Kingdom), we wrapped filters individually in aluminum-foil, then autoclaved these in a 15min-exposure fast-exhaust cycle. On the evening before the day of microbiome-harvest, we set up a custom-made flow-hood on a bench in our Conviron growth-chamber, sterilized the inside of the hood by spraying liberally with 100% ethanol, then allowed the flow of clean air to purify the inside of the hood overnight. Our custom-made hood was constructed of a large plastic tub placed on its side, with the lid cut half so that a lid-portion affixed to the tub could shield the inside of the hood from above (like a sash on a regular flow-hood), whereas the bottom half was kept open to permit access to the inside of the hood. To generate a flow of clean air through the hood, we cut a large hole into the top of the hood (i.e., one of the original sides of the tub now resting on its side) to fit into that hole the top portion of an air purifier (model HPA104 Honeywell HEPA Allergen Remover, with HEPA filter of 0.3 microns; Honeywell International Inc., Morris Plains, NJ, USA). We operated the purifier at medium flow-setting, which generated an even flow through the hood and minimized any air-vortices that could draw impure air into the hood at high flow-setting. In a pilot test, Petri-plates with PDA-medium, exposed overnight to the flow inside our hood, revealed no visible growth within seven days of incubation of these plates at room temperature. Early on a day of a between-generation microbiome-transfer, we moved the tubs with racks of planted, cold-stratified seeds from the cold-room into our growth-chamber, to have sufficient time for completion of all microbiome-harvests and -transfers (the total time needed on the day of microbiome harvest for completion of harvests/transfers of all lines was 8-10 hours, plus an additional 2 hours for distribution of pots in pre-determined randomized arrangements across 8 racks used to support pots in the growth-chamber). We began microbiome-harvests and -transfers immediately after moving pots into the growth chamber, so transferred microbiomes would interact with seeds at the very early stages of germination.

### Phenotyping of Plants; Quantification of Above-Ground Biomass

To select the two best-growing plants from a particular selection line on the day of microbiome-harvest and -transfer, we moved all eight pots from a selection line into a separate, ethanol-sterilized rack, recorded the number of leaves of each plant, and arranged plants visually by apparent above-ground biomass into a size-ranked series (Figure S3). We chose visual sizing rather than weighing for phenotyping of plants, because visual evaluation of all eight plants in a selection-line needed only about 5-10 minutes (including recording the number of leaves for all eight plants), and because microbiomes could be harvested immediately after visually identifying a particular plant for microbiome harvest without first having to cut and weigh above-ground biomass of all plants in a selection line. We harvested rhizosphere microbiomes from only those two plants within a selection line that we visually judged to have grown the largest and second-largest above-ground biomass (Figure S3).

**Figure S3.**
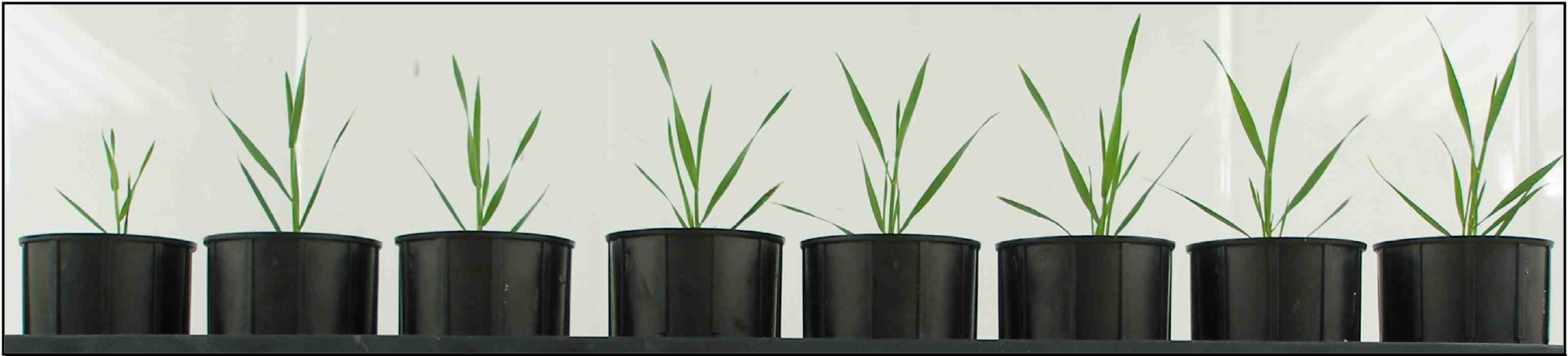
Plants of the same selection-line ranked by visually estimating above-ground biomass. The 8 plants were grown in 8 different racks (one plant per rack) in randomized positions in each rack, and the 8 plants were moved to a separate ethanol-sterilized rack for visual comparison immediately before microbiome harvesting from the two largest plants. See text for further details on *Phenotyping of Plants; Quantification of Above-Ground Biomass*. Photo by UGM.

To test the accuracy of our visual rankings, we later compared these rankings with dry above-ground (shoot) biomass. To weigh shoot-biomass, we cut each plant at soil-level at the time of microbiome harvesting, stored above-ground biomass for drying in an individual paper envelope (Coin Envelope 8cm x 14cm), dried these envelopes/plants for at least two weeks at 60°C in a drying oven, then weighed dry biomass for each plant to the nearest 0.1mg. Although we judged above-ground plant-biomass visually on the day of microbiome harvesting, of the 80 lines judged during our entire experiment (5 SOD-lines + 5 ALU-lines judged each Generation, times 8 Generations; Table S1), we picked for microbiome harvest the combination of largest (#1) and second-largest (#2) plant in 56.25% of the cases; the largest (#1) and third-largest (#3) plant in 27.50%; the largest (#1) and fourth-largest (#4) plant in 6.25%; the second-largest (#2) and third-largest (#3) in 5.00%; the second-largest (#2) and fourth-largest (#4) in 5.00%; and never any lower-ranked combination. In cases when we did not identify visually the combination of #1 and #2 plants as later determined by dry weight, the slightly lighter #3 or #4 plants were typically within 0.2-4mg (0.5-9% of total dry-weight) of the two best-growing plants in the same selection-line (Table S1). Moreover, because harvested microbiomes of the two chosen plants were mixed for propagation to the next microbiome-generation (see below *Microbiome Mixing*), we harvested in 100% of the cases the microbiomes from either the best-growing or second-best growing plant into the mixed microbiome that we then propagated to the next microbiome generation (i.e., a microbiome of one of the two best-growing plants was always included in the propagated microbiome mix). In sum, therefore, our method to visually judge plant-size was both time-efficient (about 5-10 minutes needed to visually size all plants in a selection-line and record number of leaves for each plant), ant our method was also accurate to identify those plants that had grown biomass well above-average within any given selection-line (i.e., our methods were accurate to visually identify plants that were likely associated with microbiomes that conferred salt-tolerance to plants).

In some cases, on the day of microbiome-harvest, more than two plants of the same selection line appeared to have the largest above-ground biomass. To decide between those plants for microbiome-harvest, we considered as a second criterion also the growth trajectory recorded from the day of germination to the day of microbiome-harvest, choosing then the plant with the best growth-trajectory. We quantified growth trajectory of plants during each generation with three methods: (i) measuring the length of the first leaf on Days 2-5; (ii) after Day 5, recording the number of leaves grown by a plant every other day up to a time when plants had grown about 10 leaves; and (iii) once plants had grown about 10 leaves, visual ranking of relative plant size (visual appearance of overall biomass) on a 10-point scale from 1-9, using the protocol below. ***Length of first leaf*:** After moving pots from the cold-room into the growth-chamber, the fastest-growing shoots became visible, as they pushed through the soil, after about 44 hours in the early, low-salt Generations, but growth rate was somewhat slower in the later, high-salt Generations when the first shoots became visible after 55-70 hours (Table S1). To quantify this early growth each selection-cycle, we estimated length of the first leaf during Days 2 & 3 visually without lifting the translucent lids from pots, but measured leaf-length on Days 4 & 5 to the nearest millimeter with an ethanol-sterilized ruler (millimeter scale printed on paper strip; Figure S4) held next to the growing leaf, using a different sterile paper-ruler for each plant so as not to transfer microbes between pots. In blind, repeat evaluations (Table S1), the visual sizing on Days 2 & 3 is accurate to about ±0.5mm for leaves less than 15mm tall, and accurate to about ±2mm for plants larger than 25mm. Despite the somewhat lower accuracy of the visual leaf-length estimation compared to the precise measurement with a ruler, we chose to visually size plants on Days 2 & 3 because that method allowed us to leave the pots covered with the translucent lids, thus preventing any influx of microbes when lifting a lid; plants therefore interacted only with the experimentally-transferred microbiomes for a total of 4 days without any influx of additional microbes, thus facilitating priority effects in microbiome recruitment into the initial microbiome assembled by a plant. ***Counting leaf number*:** The fastest-growing plants showed growth of a second leaf typically late on Day 5 (in the early low-salt Generations) or on Day 6 (in the later high-salt Generations). We counted the number of leaves regularly after Day 6 (Table S1), typically every other day. Because different plants with the same number of leaves differed in size of their youngest leaves (e.g., on the same day, some plants showed beginning growth of leaf #3, other plants extensive growth of leaf #3), we also recorded during these counts the relative size of each plant’s youngest leaf on a 5-point scale [see Table S1b&c: double-minus “–” (= well-below average for plants with that particular leaf number); single-minus “-” (= below average); average; single-plus “+” (= above average); double-plus “++” (= well above average)]. ***Above-ground biomass estimated on a 10-point scale ranging from 0-9*:** This third method gave the most precise estimate of above-ground biomass once plant had grown more than 10 leaves, and we used this method therefore every generation to obtain a relative measure of above-ground biomass a few days before the microbiome-harvesting. An experimenter first looked over all plants to gain an impression of the largest plants, of the average-sized plants, and of the smallest plants, then subdivided the entire range on a subjective 0-9 point-scale, with plants of average size to be scored as 4.5 on the 0-9 point-scale. Evaluating all plants rack-by rack, the experimenter scored and recorded sizes of all 200 plants in a Generation, then blindly re-scored all plants again rack-by-rack, then calculated an average between the two scores for each plant. Comparison of the first and second size-values for each plant (see Table S1b&c) showed that about 70% of the blind re-scoring were identical to the first score; and in most of the remaining 30% cases, the scores of the same plant differed by only a 1-point-value, and only in very exceptional cases (<2%) the scores differed by a 2-point value. Because of this high repeatability of this scoring method, we used this method every generation to obtain estimates of the relative above-ground biomass of each plant 1-3 days before each day of microbiome-harvesting.

**Figure S4.**
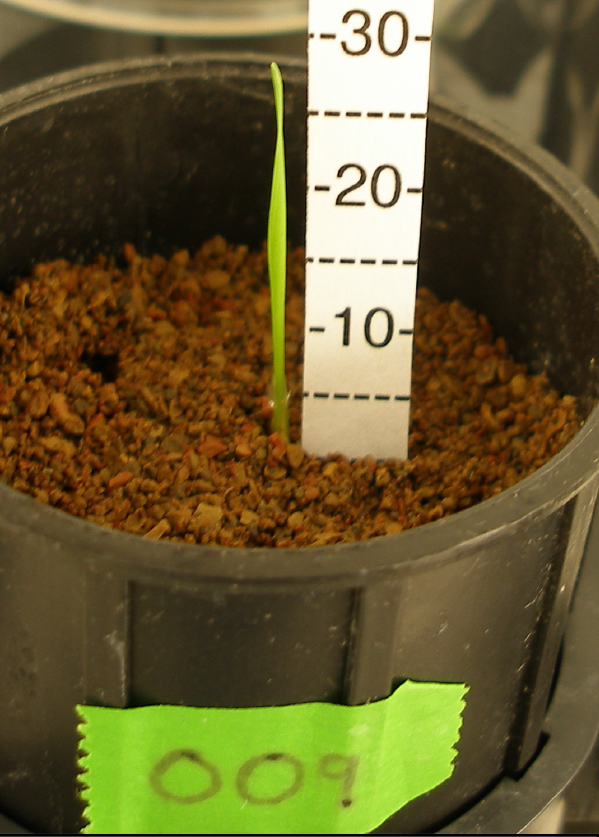
Measuring length of the first leaf on Day 3, using an ethanol-sterilized dry paper-strip with a millimeter-scale. Photo by UGM.

### Microbiome-Harvesting from a Rhizosphere & Microbiome Mixing

We performed all steps of microbiome harvest and microbiome transfer in a clean-air flow-hood (see above) set up on a bench inside our growth-chamber (i.e., we did not have to move microbiomes/pots of selection lines outside the growth chamber), and we sterilized hands and work-surfaces regularly with 100% ethanol to prevent contamination and cross-contamination of samples. After choosing the two plants with the greatest above-ground biomass (see above *Phenotyping of Plants*), we cut each plant at soil level with ethanol-sterilized scissors, stored the above-ground portion in an envelope for drying, and harvested rhizosphere microbiomes immediately to minimize microbiome changes in the absence of plant-control in the rhizosphere. To extract the root-system from a pot (Deepot) with minimal contamination, we held the shoot-stub at the soil surface with ethanol-sterilized forceps, tilted the pot such that PPC-soil would gradually loosen and fall out when squeezing the plastic pot, until the root-structure could be extracted as a whole by gentle pulling at the main root with the forceps. In most cases, the entire root structure could be extracted whole, with some loss of fine roots embedded in spilled soil. Because we were interested in harvesting microbiomes that were in close physical association with a plant (i.e., we were interested in rhizoplane bacteria, plus any endophytic bacteria if they were released during root processing as a result of any root damage), we discarded any soil adhering loosely to the roots. We dislodged loosely adhering soil by knocking the root-system gently against the wall of an autoclaved aluminum-pan (e.g., Hefty EZ Foil Roaster Pan; 32cm length x 26cm width, vertical depth 11cm) such that any dislodged soil would fall into the pan without the roots contacting any discarded soil. We then cut off the top 2 cm of the root-system (i.e., roots close to the soil surface), then transferred the remaining root-system into a 50 ml Falcon tube filled with 45 ml of salt-nutrient buffer (the same buffer used also to hydrate soils of the subsequent microbiome-generation, but diluted to half-concentration to suspend harvested microbiomes; see above *Preparations for Microbiome-Harvesting*). We repeated this process with the second plant chosen for microbiome-harvest from the same selection line, and added this second root-system to the same Falcon tube as the first root-system. Combining both root-systems for microbiome-harvesting generated a so-called mixed-microbiome collected from two ‘mother rhizospheres’ (see *Mixed Microbiome Propagation*; and Box 3 in Mueller & Sachs 2015), which we then transferred within the same selection line to all eight ‘offspring plants’ of the next microbiome-generation.

### Microbiome-Fractionation with Microfilters

To dislodge microbes from roots and from soil-particles adhering to roots, we turned a closed Falcon tube upside-down 50 times, then permitted the solids to settle in the bottom of the tube for 1 minute. A 1cm-deep sediment of PPC-soil particles typically accumulated in the bottom cone of a Falcon tube, with the roots settling on top of this sediment, and small particles and colloids remaining suspended in the salt-nutrient buffer. We aspirated 20 ml of this suspension with a sterile 20 ml syringe (external syringe diameter fitting into a 50 ml Falcon tube), then attached to the syringe’s Luer-lock a 2 μm Whatman microfilter (model Puradisc 25 GD2 Syringe Filter, 25 mm diameter; Whatman PLC, United Kingdom), then filtered the aspirated suspension into an empty sterile 50 ml Falcon tube. Making sure that the exterior of the syringe did not become contaminated during this first filtering, we repeating this step with the same syringe to filter another 15-20 ml of the suspension, then mixed the combined filtrates by inverting the Falcon tube several times. The total volume of 35-40 ml filtrate was sufficient to inoculate 8 ‘offspring plants’ each with 4 ml filtrate (total of 8 x 4 ml = 32 ml needed). In pilot tests, plating on PDA-medium 10 μL of this filtrate (2μm filter) yielded thousands of bacterial colonyforming-units (CFUs) but no fungal CFUs within 24 hours growth; whereas plating on PDA-medium 50 μL of this same filtrate that had been filtered a second time with a 0.2μm filter (VWR Sterile Syringe Filter, 0.2μm polyethersulfone membrane, 25mm diameter; Catalog #28145-501; retains even the small-sized *Bre-vundimonus diminuta*) did not yield any visible microbial growth on PDA plates kept for 7 days at room temperature. These results justified addition of a third control-treatment in Generation 9 (0.2μm filtration of suspension) to test growth-promoting effects of chemicals and viruses co-harvested with the harvested microbiomes (see below *Crossing Evolved SOD- and ALU-Microbiomes with SOD- and ALU-Stress*). Although a 0.2 μm filter may not eliminate ultra-small bacteria (e.g., Luef *et al* 2015; we did not use filters of smaller pore size because it was too difficult to press liquid through such filters with a hand-held syringe), our control comparison between 2.0 μm-filtered and 0.2 μm-filtered bacterial microbiomes can still test whether the bulk of the bacterial microbiome (in size range 0.2-2.0 μm) or alternatively any smaller-sized organisms (viruses, ultra-small bacteria) plus solutes in the soil are responsible for conferring salt-tolerance to plants in our experiment.

### Inoculation of Seeds; Transfer of Microbiomes to Plants of the Next Microbiome-Generation

During planting, the 200 pots of each microbiome-generation had been ordered numerically in the 4 racks used for stratification in the cold-room, so it was easy to locate in these racks a pot with a particular number that had been assigned to a specific selection-line and needed to be inoculated with a microbiome. To inoculate a seed planted in a particular pot, we moved the pot into our clean-hood in our growth chamber, opened the pot’s translucent cap inside the hood (using one hand to hold the pot while opening the cap with thumb and index finger of that same hand), then used a 5 ml pipetter to transfer 4 ml of the microbiome-filtrate to the center soil in a pot where a seed had been planted before vernalization/stratification. We spread the 4 ml filtrate across an area with a radius of about 5mm around a seed, applying some of the filtrate directly onto the seed (the exact location of the seed was indicated by its awn protruding above the soil; see *Planting* above), and we spread some of the filtrate also in a circular motion onto the surrounding soil within 5 mm distance of a seed. To keep the filtrate well-mixed during the time needed to inoculate all 8 ‘offspring soils’ of the same selection-line, we repeatedly mixed the filtrate in the Falcon tube with the pipette-tip before aspirating a 4 ml-aliquot to inoculate the next pot. We then taped a small tag of labeling-tape to the lid of each pot that had received an inoculum (as a check to verify later that all pots had received an inoculate, no pot was accidentally skipped, or any pot was accidentally inoculated twice), then we returned the pot to its appropriate position in one of the four racks. After inoculation of all 200 plants within a Generation, all pots were distributed among the 8 racks used to support plants in the growth chamber (see below *Randomization of Pot-Positions in Racks*).

Each pot was capped for first 4 days to promote priority effects during microbiome establishment (i.e., capping prevented immigration of extrinsic microbes into the soils/microbiomes for the first 4 days; see above *Planting*), but all caps were removed on Day4 because the tallest plants (35-40mm tall on Day4) were close to reaching the cap-ceiling. We monitored growth during first 5 days (see above *Phenotyping*) by recording length of the first leaf on Days 2-5, and recording day of appearance of the second leaf (typically on Days 6 or 7; Table S1). Seeds that did not germinate or that germinated very late (i.e., no above-ground growth visible by Day 4) were extracted from pots with forceps and inspected. Most of these seeds had failed to grow both a rootlet and shoot by Day4, but some seeds had grown a rootlet but no shoot. In a typical microbiome-generation, about 88-100% of the plants showed a visible shoot within the first 3 days (Table S1). Germination rates were therefore good overall, and most lines had the planned 8 replicates (sometimes 7 replicates, rarely 6 replicates, if some seeds failed to germinate; see Table S1). Germination-rates were often minimally higher in the Null-Control treatments compared to other treatments of the same soil-stress (slightly fewer non-germinating seeds in Null-Controls), and, across all plants, germination-rates were minimally higher in ALU-soil than in SOD-soil (see Tables S1a-c); we did not analyze these trends for statistical significance because differences were minimal, but we simply note here these general patterns became apparent only when pooling information across all 10 selection cycles.

### Randomization of Pot-Positions in Racks in Growth-Chamber

Deepots were supported in D50T racks (Stewe & Sons, Tangent, Oregon, USA). Each rack can hold a total of 50 pots (5 rows of 10 pots each), but to prevent contact of leaves from different plants and to reduce accidental between-pot transfer of microbes during watering (see below *Watering*), we used only 25 rack-positions (25 pots per rack, total of 8 racks, for a total number of 200 pots per selection cycle). Pots within a selection line were first assigned by blocking to particular rack (e.g., of the 8 replicates within a selection line, one replicate was assigned to each of the 8 racks). Within each rack, however, we randomly assigned pot positions, using the *Random Sequence Generator* option at Random.Org (www.random.org/sequences/). For Generations 0-8 (growth cycles 0-8), Tables S1a&b lists pot positions (#1-#25) from different treatments within each rack (Rack #1-8), corresponding to the following pot arrangement:

|  |  |  |  |  |  |  |  |  |  |
| --- | --- | --- | --- | --- | --- | --- | --- | --- | --- |
| #1 |  | #2 |  | #3 |  | #4 |  | #5 |  |
|  | #6 |  | #7 |  | #8 |  | #9 |  | #10 |
| #11 |  | #12 |  | #13 |  | #14 |  | #15 |  |
|  | #16 |  | #17 |  | #18 |  | #19 |  | #20 |
| #21 |  | #22 |  | #23 |  | #24 |  | #25 |  |

For the final Generation 9 when we added two more control-treatments (details below), we randomized 400 pot-positions by first assigning a pot to one of 8 racks, then randomizing position within each of the 8 racks (50 pots/rack; the position-numbering of pots shown for Generation 9 in Table S1c for each rack is numbered consecutively, starting in left top corner, without leaving empty spacer-slots between pots).

The 8 racks were positioned in two groups of 4 racks each on two comparable shelves at either side of the growth chamber. Within each selection cycle, we rotated these 8 racks in clockwise rotation each day (moving one rack from right shelf to left shelf, and one rack from left to right shelf), and at the same time we also turned each rack (such that the rack-side facing the chamber wall one day faced the chamber center the next day). This rotation-turning scheme aimed to minimize possible environmental influences dependent on location of a rack on the two shelves, and to reduce any minimal differences in light-level, air-circulation, or any such uncontrolled environmental factors that may exist between different positions on the two shelves in our growth chamber. Despite our effort to minimize rack effects through daily rack-rotation and rack-turning, as well as randomization of processing order (e.g., watering, phenotyping, microbiome-harvesting), we had occasionally racks of poorer or better plant growth (e.g., Rack 7 of Generation 9 had lower average seed production compared to other racks, because many plants in that rack did not flower, or flowered late). We do not know the exact causes for occasional small rack-effects, because we believe we treated plants across all racks equally.

### Starter Inoculum for Microbiomes at Beginning of the Experiment for Baseline Generation 0

We used a single inoculum-batch to inoculate all replicate pots of the initial baseline Generation 0. To prepare that inoculum, we filtered bacterial communities from a mix of roots and adhering soil taken from three principal sources: (a) root-systems with adhering soil of three local grass species (*Bromus* sp., *Andropogon* sp., *Eragrostis* sp.) collected into individual plastic bags on 3.Jan.2015 (about 90 minutes before microbiome harvesting) at restored native habitat at Brackenridge Field Lab of the University of Texas at Austin (www.bfl.utexas.edu/); (b) root-systems with adhering soil of 40 16-day-old *B.distachyon* Bd3-1 plants grown in PPC-soil Deepots as part of a pilot experiment quantifying the effect of salt in soil on the growth rate of *B. distachyon* (see below *Salt Treatments*); and (c) old root-systems with adhering soil of 15 Bd3-1 plants grown in PPC-soil Deepots, but that had been stored in the soil/Deepots in a cold-room (6°C) for 7 months after completion of a previous low-nutrient microbiome-selection experiment. We combined roots and rhizosphere soils from these three sources in order to capture a diversity of microbes into our starter inoculum, and we included Bd3-1 rhizospheres in order to capture specific microbial taxa that may be readily recruited by *B. distachyon* into its rhizosphere microbiomes. We suspended this mix of roots and rhizosphere soil in 200 ml e-pure water, blended the mix for 30 seconds in an autoclaved Waring blender to generate a liquid slurry, allowed the solids to settle in the blender for 1 minute, then decanted the supernatant into a separate autoclaved beaker. Adding each time 200 ml e-pure water, we repeated this blending/decanting with the remaining slurry three more times to collect a total of about 600 ml supernatant. Using vacuum filtration, we pre-filtered this supernatant in a Buchner funnel through filter paper (Ahlstrom filter paper S02-007-42), eliminating larger particles suspended in the supernatant. To harvest only bacterial microbiome components (and viruses) from this pre-filtrate, we filtered the supernatant a second time in a laminar-flow hood, using a 60 ml syringe fitted with a 2 μm Whatman™ microfilter (Puradisc 25 GD2 Syringe Filter, 25 mm diameter; Whatman PLC, United Kingdom) to generate the bacterial mix for inoculation of replicate pots of our initial baseline Generation 0. Because the Puradisc filters became clogged after filtration of about 70-100 ml supernatant, we used 8 Puradisc filters to process about 600 ml of filtrate. We reserved 500 ml of this filtrate for inoculation of 160 randomly-assigned pots in a Bacterial-Inoculate treatment (80 Bacterial-Inoculate with SOD soil, 80 Bacterial-Inoculate with ALU soil), and filtered the remaining 100 ml with 0.2 μm filters (VWR Sterile Syringe Filter, 0.2μm polyethersulfone membrane, 25 mm diameter; Catalog #28145-501) for inoculation of 40 pots in Null-Control treatments (20 Null-Control with SOD soil, 20 Null-Control with ALU soil). The Null-Control treatments controlled for, after elimination of bacteria, the effect of any chemicals and viruses that may have been co-harvested from rhizosphere roots and soils. Seeds in the Bacterial-Inoculum and the Null-Control treatments were inoculated following the procedure described above (see *Inoculation of Seeds*), except that each seed of Generation 0 received 2 ml inoculate, whereas each seed of subsequent Generations 1-9 received 4 ml inoculate transferred between generations. During inoculation of seeds, we mixed the stock filtrates regularly to prevent bacterial sedimentation and to insure standardized inoculation of all replicates in a treatment. We needed about 3 hours to complete the entire process from root collection to conclusion of all filtration steps, and another 2 hours to apply inoculate-aliquots of the filtrates to each of the assigned pots. We then moved all pots immediately into our growth chamber, and set out all 200 pots of Generation 0 into randomized positions in 8 racks (see above *Randomization of Pot Positions*; Table S1).

To test for live bacteria in our 2 μm filtrate used as the Starter Inoculum, we plated on PDA-medium (2 replicate plates) 10 μL each of the 2 μm filtrate and maintained plates at room temperature; the plates showed thousands of bacterial colony-forming-units (CFUs) within 24 hours, but no fungal growth within 7 days. To test for absence of live bacteria in our 0.2 μm filtrate, we plated on PDA-medium (3 replicate plates) 50 μL each of the 0.2 μm-filtrate; these platings did not yield any visible growth on the PDA plates kept for 7 days at room temperature. These results indicate (i) a great abundance of live bacteria (and apparently no live fungi) in our initial inoculum, and (ii) elimination by the 0.2 μm filters of live bacteria that would be apparent when plating out such filtrate on PDA plates. The latter justified our use of a third control-treatment in Generation 9 (0.2 μm filtration of suspension to test growth-promoting effects of chemicals and viruses co-propagated with the harvested bacterial microbiomes; see below).

### Selection of Microbiomes from Generation 0 to Inoculate Plants from Generation 1

At the start of our experiment, we did not assign microbiomes (i.e., pot numbers) from Generation 0 to specific selection lines, to permit selecting the best-growing plants from Generation 0 to contribute microbiomes to the selection-lines starting with Generation 1. We chose this particular assignment rule because random assignment to selection lines would result in some cases for a poorly-growing plant to contribute microbiomes to Generation 1, and we wanted to increase the chance of obtaining a response to microbiome-selection in the fewest rounds of selection. To select plants for harvesting and propagation of rhizosphere microbiomes, we ranked, separately for plants in the SOD and ALU treatments, the plants in the Bacterial-Inoculate treatments of Generation 0 by relative size, then picked the 10 best-growing plants of each salt-treatment to contribute microbiomes to the selection-lines that we started with Microbiome-Generation 1 (Table S1). On Day 22 of Generation 0 (day of microbiome harvest and microbiome transfer; Table S1), we first ranked plants by relative size-scores (i.e., average size-score averaged across three scores received by a plant on Days 18, 19, 20; see protocol *Above-Ground Biomass Estimated on a 10-Point Scale*), then used number of leaves recorded on Day 21 as a second criterion to differentiate between plants of equal average size-score. Among the 10 best-growing plants within each of the SOD and ALU salt-treatments, we paired plants randomly to generate 5 combinations (2 plants each) for mixing of harvested microbiomes within each pair (i.e., harvested root-systems were combined from the two plants to harvest a mixed microbiome from both plants, as described above for *Microbiome Mixing*). Within each of the SOD and ALU treatments, the 5 mixed microbiomes from Generation 0 were assigned randomly to 5 SOD and 5 ALU selection lines (each with 8 ‘offspring microbiome’ replicates per line) that started with Generation 1. Microbiomes were harvested and processed from chosen rhizospheres as described above. At the end of Generation 0, as well as at the end of each subsequent Generation, we cut all plants at soil level to preserve above-ground growth for later weighing of dry biomass for each plant (Table S1; see also above *Phenotyping*).

### Salt- and Control-Treatments in Generation 0-9; Sample Sizes Per Treatment

Starting with Generation 1 and continuing until the last Generation 9, we included the two aforementioned salt-treatments (SOD and ALU soil) with *5 SOD Microbiome-Selection* Lines (8 replicates each, for a total of 40 replicates) and with 5 *ALU Microbiome-Selection* Lines (8 replicates each, for a total of 40 replicates). Also starting with Generation 1 and continuing until the last Generation 9, we included two control treatments for each of the SOD and ALU treatments, *Null-Control* (on SOD- and on ALU-soils) and *Fallow-Soil Microbiome Propagation* (on SOD- and on ALU-soils). ***Control 1, Fallow-Soil Microbiome Propagation***: For this control, we harvested microbiomes from fallow soil (from pots without a plant), then propagated the harvested microbiome to sterile fallow soil to perpetuate ‘Fallow-Soil Microbiomes’ in the absence of plant influences (e.g., absence of plant secretions into the soil). Fallow-soil pots were treated throughout each selection-cycle exactly like pots with plants; for example, these fallow-soil pots received the same amount of water whenever all other pots were watered. Each Fallow-Soil-Line had only one replicate pot, so a microbiome harvested from fallow-soil was propagated to a single pot of the next selection-cycle to continue a particular Fallow-Soil-Line; a portion of the same microbiome harvested from fallow-soil was also transferred to pots with plants of the next cycle to assay the effect of a harvested fallow-soil-microbiome on plant growth (but those microbiomes were later not propagated to subsequent Generations; i.e., these inoculations of control plants aimed at assaying the effect of un-selected fallow-soil-microbiomes on plant growth under the increasing salt stress that we increased stepwise between Generations; see above *Logic of Salt-Stress Ramping*). We chose a control of fallow-soil microbiome-propagation because this treatment resembles the kind of microbiome conditions that many plants encounter in horticulture and agriculture (soils are left fallow for some time before planting). Changes in fallow-soil microbiomes between Generations reflect ecological changes as microbe communities change over time, as well as any microbial immigration from external sources (e.g., airborne microbes raining into the soil; perhaps also unintended accidental cross-contamination between soils from different pots). We initially allocated 8 control-replicate test-plants per Fallow-Soil-Line to test the effect of each harvested fallow-soil microbiomes on plant growth (total of 5×8=40 replicates for SOD, 5×8=40 replicates for ALU), but we reduced the number of control-replicate test-plants for each of the 5 Fallow-Soil-Control replicates per line in later Generations (first reducing the number to 6 control-replicate test-plants per line in Generation 4; then reducing the number to 4 control-replicate test-plants per line in Generation 5-9), because it became clear during the first few Generations that plants receiving fallow-soil-microbiomes grew poorly under the salt stresses, far inferior to plants in the corresponding selection-lines where plants received artificially selected microbiomes (i.e., we could differentiate averages between fallow-soil and microbiome-selection lines even with the smaller number of control-replicate test-plants in the fallow-soil controls). **Control 2, *Null-Control***: For this control, plants received no experimental microbial inoculation; instead, these control plants received on the day of microbiome transfer an aliquot of the same sterile salt-nutrient buffer that we used to harvest microbiomes and then transfer to seeds of the next Generation. Because our pots were capped for the first 4 days of seed germination, Null-Control-plants grow initially under sterile conditions (before caps are lifted on Day4), but airborne microbes can enter the sterile soil and rhizospheres of Null plants from the air after Day4 once caps are lifted from pots. In pilot experiments, Null-Control plants invariably grew better during the first 10-20 days than any plant inoculated with microbiomes (see Table S1a-c), possibly because Null-Control plants do not need to expend resources to mediate interactions with microbes, or because Null-Control plants do not have to compete with microbes for nutrients in the soil. Despite the microbially unusual soils of Null-Control plants, we included this control treatment because it was easy to set up (no microbiomes needed to be harvested to inoculate Null-Control soils), because Null-Control conditions were easy to standardize within Generations, and because Null-control Conditions may even be standardized between Generations if microbial immigration (i.e., rain of airborne microbes) into Null-Control soils can be assumed to be relatively constant over time. We initially allocated 10 replicates of SOD pots and 10 replicates of ALU pots to Null-Controls, but we increased the number of replicates in later Generations for the Null-Control treatments (first we increased to 20 replicates in Generation 4, then to 30 replicates in subsequent Generations) in order to improve the estimates (reduce confidence intervals) of the average growth of plants in Null-Control treatments. Sample sizes for all treatments are summarized for all Generations in Table S1a-c.

### Watering During Each Selection Cycle

We watered pots such that the total weight (pot plus hydrated soil) stayed between 200-250g and did not exceed 260g. We found in pilot experiments that a pot would be over-hydrated if the total weight reached 260-270g or more, which would result in dripping of excess water from the bottom of the pot, thus leaching nutrients and salts. Keeping pot weights well below 260g at all times therefore prevented leaching of nutrients and salt. To prevent cross-contamination (microbe-exchange) between pots, we did not use bottom-hydration by immersing racks in a waterbath, but we watered pots individually, only from above, and always with autoclaved water that we dispensed with a Seripetter Dispenser (adjustable to dispense volumes of 2.5-25ml; BrandTech Scientific Inc; Essex, CT, USA) mounted on a 6-liter carboy. Because we kept pots capped during the first 4 days of plant growth (we removed caps during the afternoon of Day 4), soils remained well-hydrated during germination (little water evaporated from soil, humidity inside the cap was near 100%). We watered for the first time on Day 5 of each selection-cycle, and thereafter approximately every 2 days (sometimes also at 1-day or 3-day intervals, depending on humidity in the growth chamber and on experimenter time-constraints), but we did not preplan to follow a rigorous 2-day watering schedule (see Table S3). We typically watered 15-25 ml per pot depending on water loss, which depended on humidity in the growth chamber and on the size of plants (humidity was greatest during Generations 4&5 because of unusually high rainfall in spring 2015). To determine the volume to be watered on a given day, we selected six pots haphazardly from the 4 racks, and weighed these on a scale (sterilizing the surface of the scale with 100% ethanol before placing a pot onto the scale). The difference between the average weight of these six pots and 255mg was the maximum quantity of water to be added to each pot. The amount to be watered could be varied to the nearest 0.5 milliliter with the carboy-mounted Seripetter Dispenser. To prepare carboys, we filled each with 6 liter of e-pure water, and autoclaved the water to ensure sterile watering. Immediately before watering, we quickly opened a carboy to add a specific volume of 1-Molar salt solution to generate a desired salt-concentration in the water (details in Table S3), mixed the contents by vigorous shaking of the carboy, mounted the ethanol-sterilized Seripetter Dispenser onto the carboy, flushed the dispenser five times to eliminate any ethanol in the dispenser, then began the watering. During the days when the Seripetter Dispenser was not used, we mounted it on a 1-Liter bottle with 100% ethanol, and kept the entire dispenser filled with ethanol to prevent growth of microbial biofilms inside the dispenser. We used different carboys dedicated to watering of SOD-salt and ALU-salt, to minimize cross-contamination of salts between treatments. In each round of watering, we first watered all pots of the SOD-treatment, then rinsed the dispenser with 100% ethanol, then watering all pots of the ALU-treatment. To minimize the chance of accidentally adding the wrong salt-water to a pot (e.g., accidentally watering ALU-soil with SOD-water, or vice versa), we labeled pots of the different salt treatments with different colors (white-label for pots ##001-100 to indicate SOD-treatment, and green-label for pots ##101-200 to indicate ALU-treatment). Table S3 summarizes the exact watering schedules, volumes of water added, and salt concentrations of the water added.

**Table S3.**
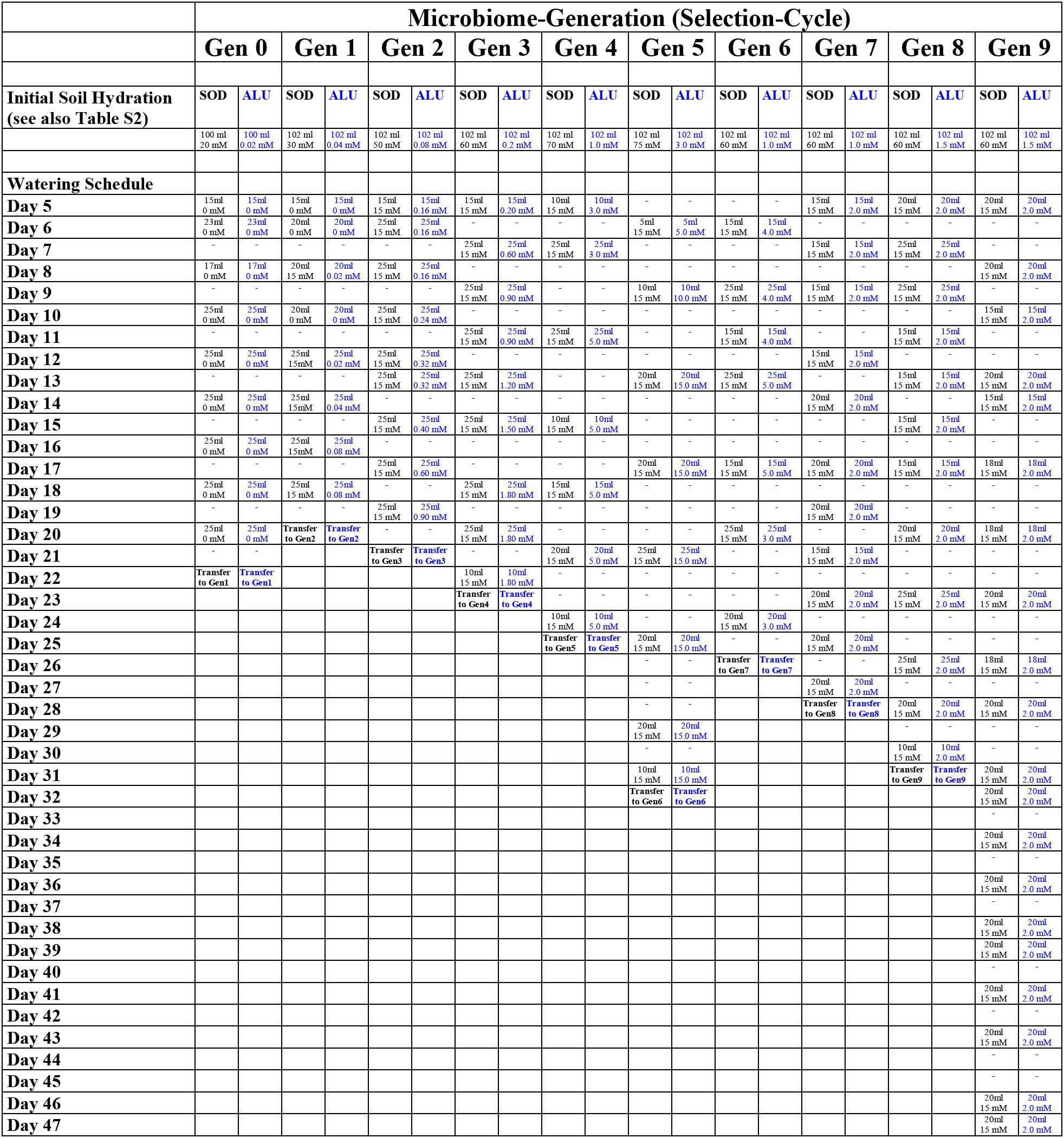
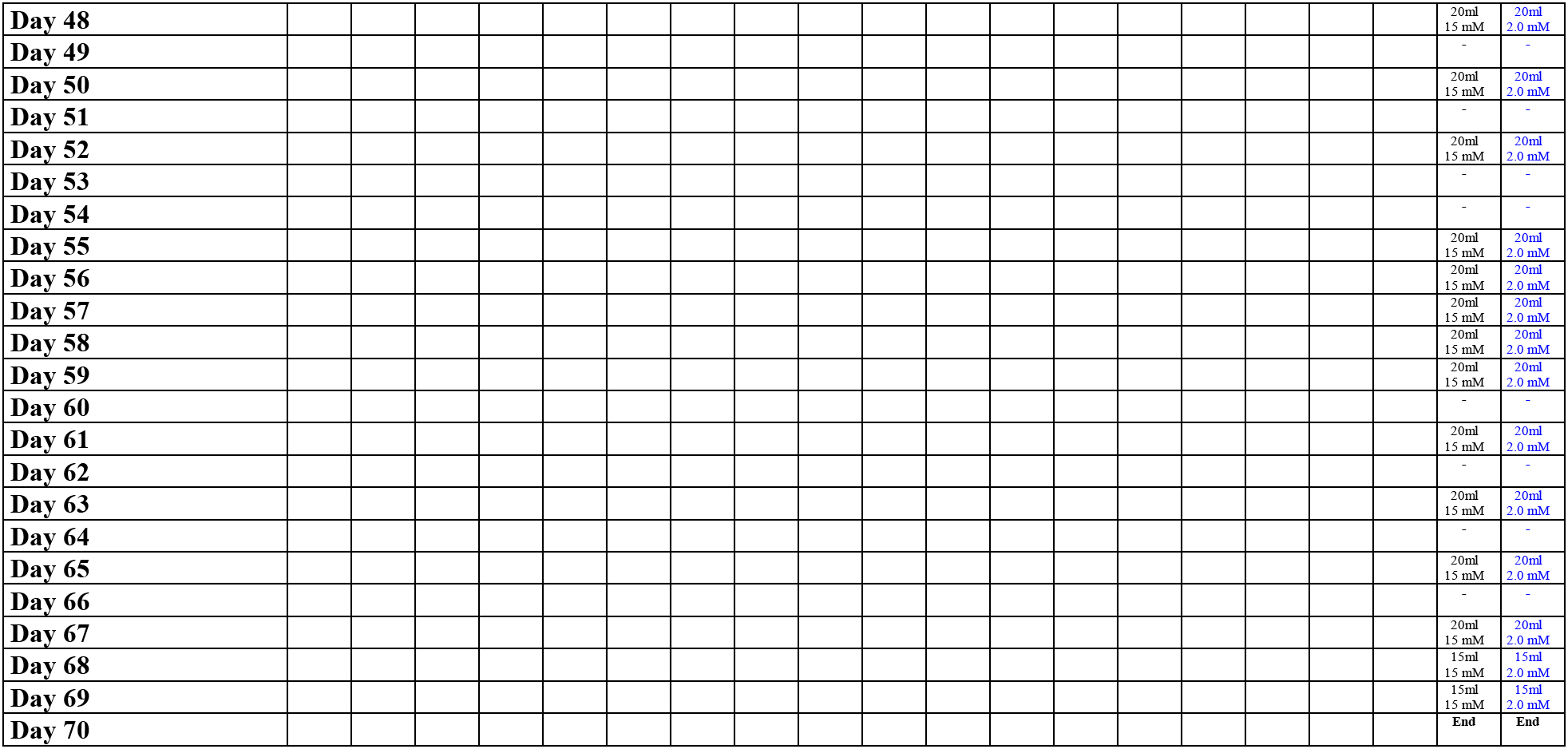
Ramping Salt-Stress Between and Within Microbiome-Generations. Soil in each pot was initially hydrated with 102 ml salt-solution [94 ml added to soil prior to autoclaving; 4 ml during planting; and 4 ml during microbiome inoculation in Generations 1-9 (2 ml in Generation 0); see *Planting* and *Inoculation of Seeds*]. In the baseline Generation 0, plants were watered only with unsalted water, but starting with Generation 1, we increased salt-stress *within* each Generation by watering with salted water. Because we capped pots for the first 4 days to control initial microbiome assembly, we started watering each Generation on Days 5 or 6. Pots of Generations 0-3 were watered more because of low humidity (because of heating of Greenhouse Facility in winter) and pots of Generations 4&5 were watered less because of high humidity (unusual rainfall in spring, increasing general humidity; see *Growth Chamber*). Pots were watered more in the second half of Generation 9 because plants grew large and transpired more water. Plants of Generations 0-8 were short-cycled to grow only for 20-30 days before microbiome transfer (to about 10-15 leaves for the largest plants); plants in Generation 9 were grown for 68 days to permit ripening of seeds. SOD = sodium-sulfate; ALU = aluminum-sulfate.

### Flowering

Because we short-cycled plants in Generations 0-8 and harvested microbiomes when plants were relatively young (20-30 days old; the largest plants had typically 10-15 leaves), only few plants bolted and developed flowers during the short-cycled Generations 0-8; all these cases of flowering were in Generations 1 & 8 (Table S1a&b), whereas no plants flowered in Generation 0 and in Generations 2-7. The long light-phase (20h light, 4h dark) stimulated flowering uniformly in each Generation, but our short-cycling scheme aimed to harvest microbiomes well before plants began to flower. Because of scheduling-constraints, Generation 8 was grown for slightly longer (31 days) than earlier Generations, which could explain the flowering in some of these plants, but it is unclear why some plants began to flower in the far shorter Generation 1 (20 days). Plants in Generation 9 were grown for 68 days to permit seeds to ripen, and most of these plants flowered (Table S1c). The fact that not all plants flowered in Generation 9, and the observation that onset of flowering was delayed in the control-treatments (Table S1c), indicate that plants were indeed stressed by the salts, because in salt-free soils virtually all plants would have flowered.

### Analysis of Relative Plant Performance

Because successive generations were not grown under precisely identical conditions (e.g., we had to increase the duration of selection-cycles in later generations because plant-growth decelerated slightly under the increasing salt-stress; we had to adjust watering schedules because of uncontrolled humidity in our growth chamber), we plot in Figure 1 plant-performance as relative above-ground biomass (rather than absolute biomass), relativizing the observed dry-biomass of a specific plant by average biomass across all plants in that plant’s salt treatment (i.e., biomass of a SOD-plant is relativized with respect to average biomass across all SOD-plants; likewise for ALU-plants). The overall average across all plants within a salt-treatment in a given Generation is therefore 1, and plants (and selection lines) performing poorer than the average have scores <1, whereas plants or lines performing above the average have scores >1 (see Figure 1). To calculate relative plant performance, we used the dry-biomass grown by a plant during its selection-cycle, as summarized in Figure 1 and in Tables S1a-c. Records of absolute above-ground dry-biomass for all plants of all Generations are also given in Tables S1a-c. In Generation 9, we allowed plants to grow for 68 days to permit ripening of seeds; plant performance and meta-genomic information of plants in Generation 9 are therefore not comparable to previous Generations, but treatments within Generation 9 can be compared.

### Crossing Evolved SOD- and ALU-Microbiomes with SOD- and ALU-Stress in Generation 9; Solute-Control in Generation 9

At the end of the experiment in Microbiome Generation 9, we modified protocols in three important ways: (a) we grew plants for 68 days to permit flowering and ripening of seeds, because seed production seemed a more informative estimator of plant fitness than the proxy of above-ground biomass used in earlier Generations; and (b) we doubled the total number of pots to 400 (i.e., 400 plants) to permit addition of two control treatments (in addition to Fallow-Soil and Null-Control treatments already used in earlier Generations). We added these two control treatments to understand the mechanistic basis of the salt-tolerance-conferring effects of microbiomes in the SOD and ALU selection lines. The first additional control was *Solute Control*, where we filtered out all live cells from the harvested microbiomes in the selection lines (using a 0.2μm filter; see above *Microbiome-Fractionation with Microfilters*), to test the growth-enhancing effect of plant-secreted solutes and viruses that may be co-harvested and co-transferred with microbiomes in the selection-lines. The second control was 2×2 *Cross-Fostering Control* where we crossed harvested microbiomes from the SOD and ALU selection lines with the two types of salt stress in soil (i.e., microbiomes harvested from SOD-selection-lines were tested in both SOD-soil and in ALU-soil; microbiomes harvested from ALU-selection-lines were tested in both SOD-soil and in ALU-soil) to test specificity of the salt-tolerance-conferring effects of the microbiomes. This Cross-Fostering treatment allowed us to address the question whether the salt-tolerance-conferring effects of the *SOD-selected* microbiomes confer these effects only under SOD-stress, or also in ALU-stress; and vice versa the additional question whether the salt-tolerance-conferring effects of the *ALU-selected* microbiomes confer these effects only under SOD-stress, or also in ALU-stress. This basic cross-fostering design was inspired by the experimental methods developed by Lau & Lennon (2012), except that, in contrast to Lau & Lennon (2012), our plant-populations did not evolve, and that we artificially selected on microbiomes (whereas in Lau & Lennon plant populations evolved under artificial selection and microbiomes were allowed to change ecologically, without artificial microbiome selection).

### Phenotyping of Plants and Microbiome-Harvesting in the Last Generation 9

In contrast to Generations 0-8 when we used early growth of plants (above-ground biomass during first 3-4 weeks) as host-phenotype to select indirectly on microbiomes, in the last Generation 9, we allowed plants to mature for 68 days (10 weeks), such that plants could flower and seeds could ripen. Because of the longer growth, some plants started to senesce towards the end of Generation 9 and some individual flower-stalks of some plants started to dry (no plant dried completely by the end of Generation 9); comparisons of metagenomic information from Generation 9 versus earlier Generations therefore need to be interpreted with caution (i.e., metagenomic information from Generation 9 is best compared between treatments within that Generation). Despite these limits of metagenomic comparisons pertaining to Generation 9, we decided to grow plants to seed in this last Generation because we were interested in understanding how treatment differences in above-ground biomass apparent at Days 20-30 (when we harvested microbiomes in Generations 0-8) would translate into treatment differences in flowering and seed-set if plants were allowed to grow longer. Apart from the longer duration of Generation 9 to permit flowering, a second important difference is likely the gradually increasing salt-concentration in soils of Generation 9 that were watered 34 times with salted water over 68 days (Table S3), in contrast to watering with salted water fewer times over the shorter 20-30 days in Generations 0-8 (9-12 waterings, depending on the Generation; see Table S3). At the end of Generation 9, all plants were cut at soil level, above-ground biomass was preserved for each plant in individual envelopes (for drying and later weighing of seeds and overall biomass; see above), and each root systems was extracted from its pot and placed into an autoclaved aluminum-tub for further processing. Harvested root-systems of plants from Generations 0-8 were comparatively small (filling about 30-60% of the soil-volume in each pot), but root-systems at the end of Generation 9 were large and extended through the entire soil-volume in each pot. We shook-off most of the adhering soil from each root system of Generation 9, cut off and discarded the top-most 2cm portion with sterile scissors, then cut the remaining root-system lengthwise (top to bottom) to preserve half of the root-system in 100% ethanol (for metagenomic screens of bacterial communities), whereas we flash-froze (in liquid nitrogen) the other half of the root-system for possible later transcriptomics analyses. For some of the best-growing plants in the SOD- and ALU-selection-lines, we also preserved a representative portion of the root-system in sterile 20% glycerol (for storage at −80°C for possible later isolation of microbes). Processing all root-systems (nearly 400 plants) took considerable time over three successive days (Days 68-70 of Generation 9). Although we processed plants from 3 racks on Day 68 (Racks #3, #8, #7), 3 racks on Day 69 (Racks #6, #2, #4), and 2 racks on Day 70 (Racks #1, #5), we summarize all weight-data of these plants in Table S1c in columns labeled Day 68.

### Statistical Analyses *Plant Biomass, Generations 1-8*

We performed all analyses in R v3.3.1. We assessed differences in above-ground plant biomass (dry weight) among treatments of Generations 1-8 by fitting the data to a generalized linear mixed model with a gamma error distribution. Line was entered as a random effect; generation, treatment, and their interaction were entered as fixed effects. Statistical significance of fixed effects in the GLMMs were assessed with likelihood ratio tests and Tukey tests employed for post-host comparisons of treatment means. Selection of the appropriate error distribution for the GLMMs was evaluated by visual inspection of Q-Q plots, and homoscedasticity was assessed using plots of the residuals of the model against the fitted values. Because plants were short-cycled in Generations 1-8 (i.e., grown long enough so plants produce typically 9-15 leaves, too short to bolt and produce flowers; see Table S2), plants did not produce any seeds, and therefore only above-ground plant biomass (dry weight) could be compared between treatments of Generations 1-8.

### Statistical Analyses: *Total Seed Weight, Generation 9*

Because plants were grown long enough to flower in Generation 9, we compared total seed weight per plant among microbiome-selection treatments (plant present; microbiomes are harvested from plants for transfer to seeds), Fallow-Soil Control (no plant present; microbiomes are harvested from fallow soil for transfer to seeds), and Null-Control (no initial microbiome inoculation, microbes establish in microbiomes when microbes “rain in” from the air). Because plants were strongly salt-stressed in Generation 9 and many plants therefore did not flower or only produced very few seeds, the distribution of data was not normal (Figure S5 top-left). We therefore attempted several data-transformations to approximate normality, including *square-root(seed weight)* transformation (Figure S5 top-right), *log(seed weight)* transformation [excluding the plants that generated zero seeds because log(0) is undefined; Figure S5 bottom-left], and *log(seed weight + 1)* transformation (making it possible to retain the plants that produced zero seeds, because seed-weight values of all plants was increased by 1mg; Figure S5 bottom-right). None of these transformations generated a distribution that approximated normality (Figures S5b-d). We therefore used Kruskal-Wallis tests for non-parametric evaluation of treatment differences; and we used Mann-Whitney U-tests for non-parametric post-hoc comparisons between treatment means, correcting p-values using the false discovery rate. All tests were two-tailed with alpha=0.05.

**Figure S5.**
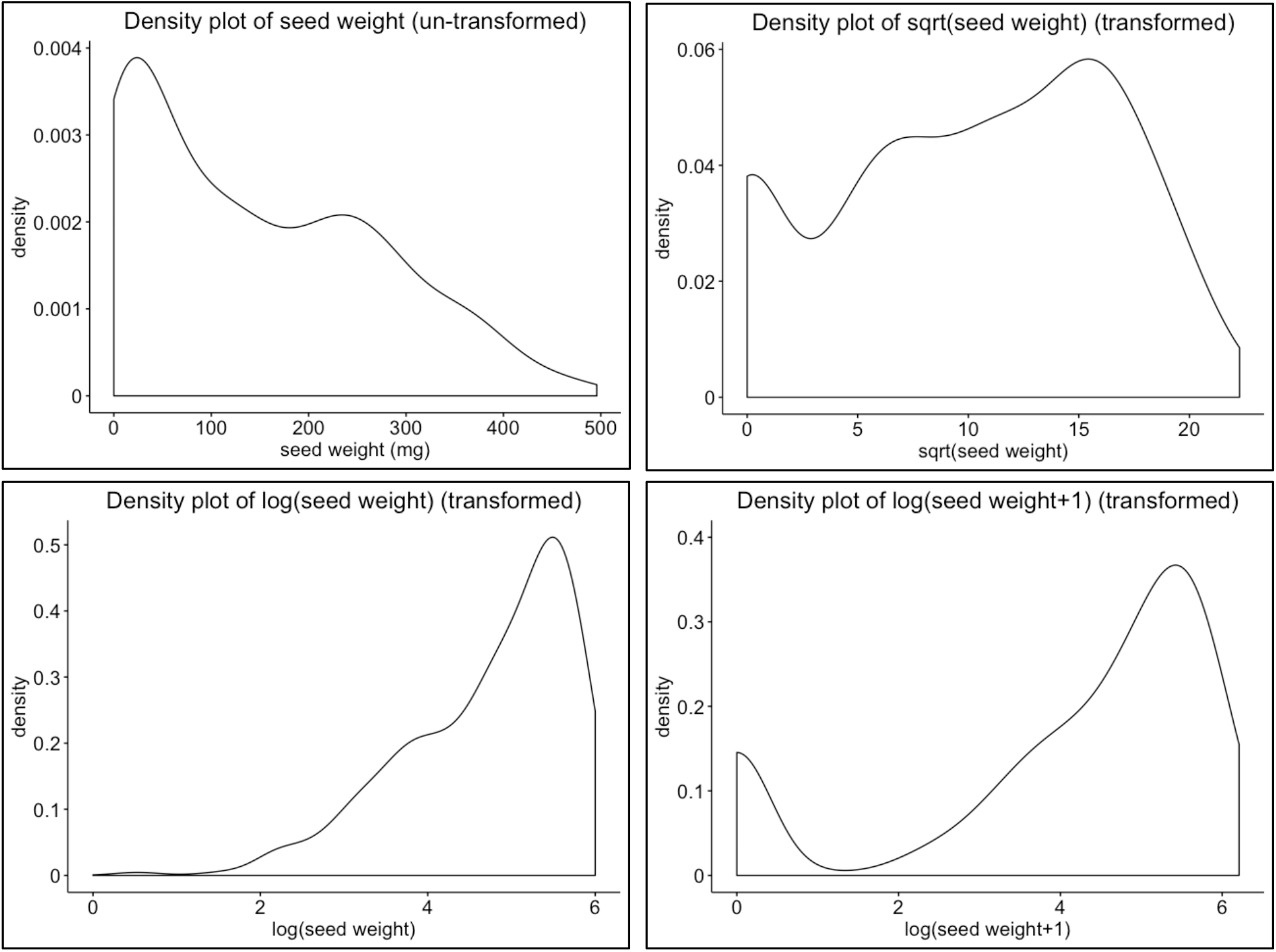
**Top-left:** untransformed seed-weight data in milligram (mg), indicating a skewed distribution, with many plants producing no or few seeds because of the extreme salt stress during Generation 9. **Top-right:** square-root transformed seedweight data. **Bottom-left:** log-transformed seed-weight data, excluding seed-weights of zero because log(0) is not defined. **Bottom-right:** log-transformed seed-weight+1 data. None of the three transformations generated a distribution that approximated normality, and we therefore used non-parametric tests to evaluate differences in seed production between treatments.

## RESULTS

### Generations 1-8: Effects of differential microbiome propagation under sodium-sulfate (SOD) stress

We found a significant main effect of treatment on plant biomass over 8 generations of microbiome selection under sodium-sulfate stress (LRT: Treatment, Chisq=27.8, p<0.001; Generation, Chisq=381.8, p<0.001; Treatment x Generation, Chisq=15.2, p=0.37; Figure 2 left). Plant biomass was 75% higher in the plant-present microbiome-selection lines (beta=0.57 ± 0.06, z=10.0, p<0.001) than in the fallow-soil control lines, and 66% higher than in the null-control line (beta=0.50 ± 0.07, z=7.4, p<0.001). There was no significant difference in biomass between the fallow-soil and the null-control treatments (beta=0.07 ± 0.06, z=1.1, p=0.29). The lack of a significant interaction between treatment and generation (Chisq=15.2, p=0.37) indicates that gains in plant biomass were realized quickly in the first few selection cycles, and that the advantage of the plant-present treatment over the fallow-soil treatment was maintained as the concentration of sodium-sulfate was ramped up over the course of the experiment.

### Generation 9, SOD-treatments

We measured total seed weight in the final Generation 9 of the experiment and found significant difference among treatments (Kruskal-Wallis Chisq=10.6, p=0.01; Figure 2 right). Total seed weight in the plant-present microbiome selection lines were 168% greater compared to the null-control line, 120% greater than the fallow-soil control lines, and 205% greater than plants grown in soil that was inoculated with filtrate (0.2μm filter) from the soil of plant-present microbiome-selection lines (Figure 2 right; Table S3).

**Table S3.** Mann-Whitney pairwise comparisons of total seed weight in the sodium-sulfate (SOD) treatments. Values represent the test statistics (p-value in parentheses) for each comparison. Significant comparisons are indicated in bold. Np = Fallow soil microbiome-propagation control, Null=Null control line, Pp=Plant-present microbiome-selection line, PpFilt=Plant-present microbiome-selection line filtrate.

|  | Np | Null | Pp |
| --- | --- | --- | --- |
| Null | 100 (0.50) |  |  |
| Pp | **0 (0.02)** | **20 (0.02)** |  |
| PpFilt | 20 (0.50) | 90 (0.71) | **20 (0.02)** |

### Generations 1-8: Effects of microbiome propagation under aluminum-sulfate (ALU) stress

Unlike the sodium-sulfate experiment, we found a significant interaction between treatment and generation under aluminum-sulfate stress (LRT: Treatment, Chisq=25.7, p<0.001; Generation, Chisq=753.7, p<0.001; Treatment x Generation, Chisq=26.6, p=0.02). The interaction was due to a drop in plant biomass in the fallow-soil treatment in Generations 4 and 5 (Figure 2). To calculate a conservative estimate of the effect size of our treatments on plant biomass, we re-ran the analysis excluding Generations 4 and 5, which eliminated the significant interaction between treatment and generation (LRT: Treatment, Chisq=17.8, p<0.001; Generation, Chisq=614.5, p<0.001, Treatment x Generation, Chisq=7.67, p=0.66). In the reduced dataset, we found that plant biomass in plant-present microbiome selection lines were 38% larger than in fallow-soil lines (beta=0.32 + 0.04, z=8.9, p<0.001), but not significantly different from the null-control line (beta=0.09 + 0.4, z=2.3, p=0.06). Null-control plants generated 26% greater biomass than fallow-soil plants (beta=0.23 + 0.04, z=5.1, p<0.001).

### Generation 9, ALU-treatments

As in the sodium sulfate experiment, total seed weight in the final Generation 9 was significantly different among treatments (Kruskal-Wallis: Chisq=9, p=0.02; Figure 2 right). Total seeds weight in the plant-present microbiome selection lines were 194% greater than in the fallow-soil lines, 101% greater than in the null-control line, and 55.4% greater than in the filtrate lines (Table S4). Plants with filtrate-inoculated soil produced total seed weights that were 89.2% greater than plants grown in the fallow-soil control (Figure 2 right; Table S4).

**Table S4.**
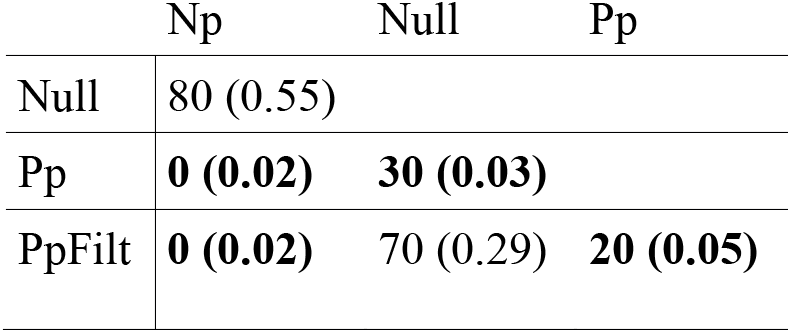
Mann-Whitney pairwise comparisons of total seed weight in the aluminum-sulfate (ALU) treatments. Values represent the test statistics (p-value in parentheses) for each comparison. Significant comparisons are indicated in bold. Np = Fallow soil microbiome-propagation control, Null=Null control line, Pp=Plant-present microbiome-selection line, PpFilt=Plant-present microbiome-selection line filtrate.

|  | Np | Null | Pp |
| --- | --- | --- | --- |
| Null | 80 (0.55) |  |  |
| Pp | **0 (0.02)** | **30 (0.03)** |  |
| PpFilt | **0 (0.02)** | 70 (0.29) | **20 (0.05)** |

### Interactions between selection history and salt stress on plant fitness

By growing plants with microbiomes from selection lines under both sodium- and aluminum-sulfate stress (*Cross-Fostering Control*), we examined whether microbiome selection produced microbiomes that conferred a salt-specific effect on plants (e.g., whether microbiomes selected to confer tolerance to SOD conferred such tolerance only under SOD stress but not to ALU stress), or alternatively whether selected microbiomes produced a generalized improvement in plant fitness for both SOD and ALU stress. There was a significant interaction between selection history and the type of salt stress to which plants were exposed in the last generation on seed mass (Analysis of deviance: Selection history, F_1,8_=<0.01, p=0.99; Salt exposure, F_1,141_=5.82, p=0.017; Selection history x Salt exposure, F_1,141_=6.42, p=0.012; Figure 2 right), indicating that performance under SOD-stress or ALU-stress in Generation 9 depended upon which salt the microbiome was selected on during Generations 0-8.

We conducted post-hoc comparisons of the treatment means and found that plants grown with microbiomes selected in sodium sulfate had total seed weights that were 70.1% greater when exposed to sodium-sulfate stress compared to exposure of aluminum-sulfate stress in Generation 9 (beta=108 ± 31.0, z=3.5, p=0.002). In contrast, plants grown in microbiomes selected in aluminum-sulfate did not differ in similar total seed weight, regardless of whether they were exposed to sodium- or aluminum-sulfate in Generation 9 (beta=4.2 ± 31.8, z=0.13, p=0.99). The effect of exposure to different kinds of salt stress on plant fitness thus depends upon the selection history of the soil microbiome.

Unlike total seed weight, there was no interaction between selection history and the type of salt stress on total plant biomass, however there was a trend toward plants growing larger under ALU-stress compared to SOD-stress irrespective of the selection history (Analysis of deviance: Selection history, F=0.14, p=0.72; Salt exposure, F=3.71, p=0.056; Selection history x Salt exposure, F=1.38, p=0.24).

